# Bystander activation across a TAD boundary supports a cohesin-dependent hub-model for enhancer function

**DOI:** 10.1101/2024.11.01.621524

**Authors:** Iain Williamson, Katy A. Graham, Hannes Becher, Robert E. Hill, Wendy A. Bickmore, Laura A. Lettice

## Abstract

Enhancers in the mammalian genome are able to control their target genes over very large genomic distances, often across intervening genes. Yet the spatial and temporal specificity of developmental gene regulation would seem to demand that enhancers are constrained so that they only activate the correct target gene. The sculpting of three-dimensional chromosome organization, especially that brought about through cohesin-dependent loop extrusion, is thought to be important for facilitating and constraining the action of enhancers. In particular, the boundaries of topologically associating domains (TADs) are thought to delimit regulatory landscapes and prevent enhancers acting on genes close in the linear genome, but located in adjacent TADs. However, there are some examples where enhancers appear to act across TAD boundaries. In these cases it was not determined whether an enhancer can simultaneously activate transcription at genes in its own TAD and in an adjacent TAD. Here, using a combination of mouse developmental genetics, and synthetic activators in stem cells, we show that some Shh enhancers can activate transcription simultaneously, not only of *Shh* but also at a gene *Mnx1* located in an adjacent TAD. This occurs in the context of a chromatin configuration that maintains both genes and the enhancers close together and is influenced by cohesin. To the best of our knowledge this is the first report of two endogenous mammalian genes transcribed simultanously under the control of the same enhancer, and across a TAD boundary. Our data have implications for understanding the design rules of gene regulatory landscapes, and are most consistent with a transcription hub model of enhancer-promoter communication.

## Introduction

How enhancers can act over very long genomic distances is still not understood. However, in mammals the correlation between the extent of regulatory landscapes of developmental genes – including that of *Shh* - and topologically associating domains (TADs) is strongly suggestive of a link to 3D genome organisation, particularly that mediated by cohesin-dependent loop extrusion (Symmons et al., 2014, 2016, Andrey and Mundlos, 2017). Indeed, cohesin is required for enhancers to activate target genes located far away in the linear genome (Calderon et al., 2022, Kane et al., 2022, Rinzema et al., 2022). A similar correlation is found between TADs and their boundaries and the ability of endogenous enhancers to activate a reporter gene integrated at different positions in the genome, suggesting that TADs confine the regulatory influence of enhancers (Symmons et al., 2014). However, there are examples in mammalian genomes where enhancers appear to be able to act across TAD boundaries (Beccari et al., 2021; Hung et al., 2023), suggesting that TAD boundaries are not completely impervious to regulatory information.

Cohesin movement along DNA in vitro can be impeded by CTCF (Davidson et al., 2023) and cohesin accumulates at oriented CTCF sites at TAD boundaries in vivo (Rao et al., 2014). However, depletion of CTCF has only a small effect on transcription in cell culture (Nora et al., 2017; Hsieh et al., 2022). Similarly, experiments to test the in vivo effects of deleting specific CTCF sites at TAD boundaries have not given clear insight into the role of TAD boundaries on enhancer function. Deletion that encompass CTCFs sites 1Mb from *PITX2* in families with a cardiac disorder result in TAD fusion and dysregulation of *PITX2* in heart tissues (Baudic et al., 2024). Deletion of all CTCF sites at a TAD boundary between the *Sox9* and *Kcnj2* loci in mice resulted in some loss of insulation between the two TADs, which was further exacerbated by deletion of additional intra-TAD CTCF sites eventually resulting fusion of the two TADs (Despang et al., 2019). However, there was no detectable change in the developmental pattern of *Sox9* expression and no overt phenotype in the animals. Similarly, deletion of individual CTCF sites at the *Shh* locus TAD boundaries did not results in dysregulated *Shh* expression or developmental phenotypes in mice (Paliou et al., 2019; Williamson et al., 2019). There is some evidence that the ability to activate across topological boundaries may be enhancer specific. At the *Sox2* locus, some enhancers are still able to activate *Sox2* in differentiation and development across inserted topological boundaries created by the insertion of arrays of CTCF sites, whereas the ability of other enhancers to activate the same gene during development is impeded by these same CTCF insertions (Chakraborty et al., 2023). Conversely, in a cancer cell line removal of all four CTCF sites at a TAD boundary led to ectopic activation of FGF3 from an enhancer in the adjacent TAD (Kim et al., 2024).

*Shh* expression in the zone of polarising activity (ZPA) of the distal posterior mesenchyme of the developing mouse limb is driven by the ZRS enhancer – located within intron 5 of the ubiquitously expressed *Lmbr1* at the opposite end of the 900kb Shh TAD from *Shh* itself (Lettice et al., 2003; Sagai et al., 2005) (Fig. 1a). However, we have previously identified that ZRS can also drive low level expression of *Mnx1*, located 150kb away in the adjacent TAD, in the developing limb bud (Williamson et al., 2019). *Mnx1* is primarily expressed in developing motor neurons of the neural tube, driven by proximal enhancers, and Mnx1 function is required to specify motor neuron identity (Arber et al., 1999; Wichterle et al., 2002; Nakano et al., 2005). However, consistent with in situ hybridisation data (Rock et al, 2007; Williamson et al., 2019) a transgene inserted at the *Mnx1* locus gives detectable lacZ staining in the ZPA of the E11.5 limb (Arber et al.,1999) and a lacZ reporter inserted 24kb upstream of the *Lmbr1* promoter, just beyond the Shh TAD boundary, shows evidence of weak expression driven by ZRS in the limb, and by Mnx1 enhancers in the the neural tube (Anderson et al., 2014). There is no reported defect in limb development in *Mnx1* knockout mice suggesting that low level *Mnx1* expression in limb development is bystander activation driven by the ZRS. This is all the more remarkable because this activation is in animals across an intact TAD boundary.

**Figure 1.**
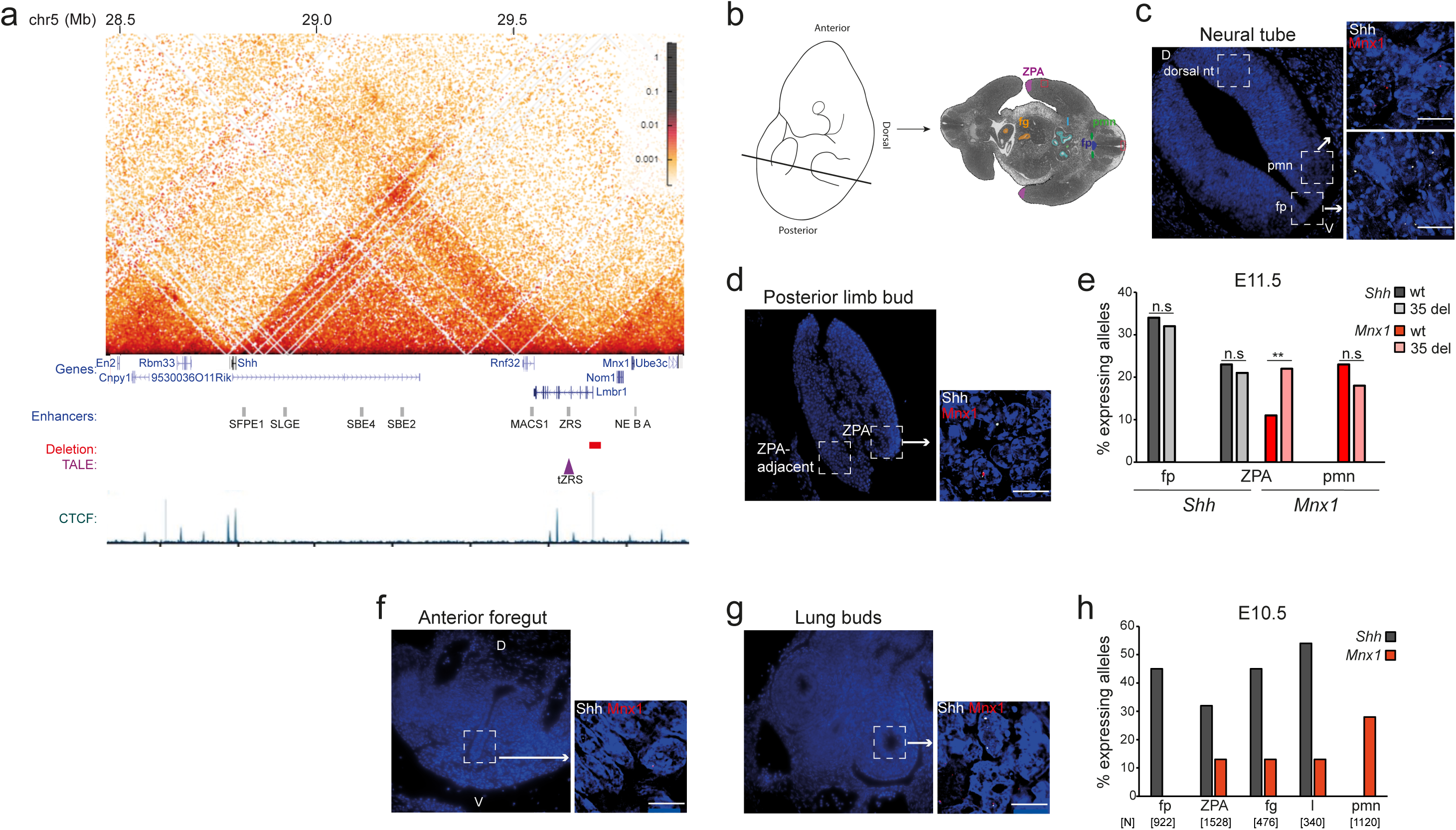
Long-range *Shh* enhancers can activate transcription at *Mnx1* in Shh expressing tissues. **a,** Hi-C heatmap, created using HiGlass, of the *Shh* TAD from wild-type mESCs, at 16-kb resolution. Data are from Boyle et al., 2020. Genes, *Shh* and Mnx enhancers, 35 kb deletion, position of the TALE target and the CTCF ChIP–seq track, are shown below the heatmap. Genome coordinates (Mb): mm9 assembly of the mouse genome. **b.** Left, mouse embryo cartoon indicating the position and orientation of the tissue sections analysed represented on the right. Highlighted are: zone of polarizing activity (ZPA; purple), floorplate (fp; blue), pre-motor neurons (pmn; green), foregut (fg; orange), lung buds (l; light blue), and non-expressing limb and neural tube tissue (red boxes). **c & d.** Representative images of tissues (left) and nuclei (right) showing RNA-FISHsignal at *Shh* (white) and *Mnx1* (red) in the (**c**) floorplate (fp) and pre-motor neurons (pmn) of the neural tube and (**d**) ZPA of the posterior forelimb bud. D=dorsal, V= ventral. Scale bars, 5 μm. **e.** % of alleles with *Shh* and *Mnx1* RNA-FISH signal in wild type and 35 kb deletion (35 del) E11.5 mouse embryos in expressing tissues of the neural tube (floorplate (fp), *Shh*, and pre-motor neurons (pmn), *Mnx1*) and the posterior limb bud (ZPA, both genes). The data were compared using a two-sided Fisher’s exact test; n.s., not significant; **, *P* < 0.01. Data for a biological replicate are in Extended Data Figure 1a. Number of alleles scored, proportions transcribed, and statistical data are in Extended DataTable 1. **f & g.** Representative images of tissues and nuclei showing RNA-FISH signal for *Shh* (white) and *Mnx1* (red) in the ventral foregut (**f**) and lung bud (**g**). Scale bars, 5 μm. **h.** The percentage of *Shh*-expressing alleles in the floorplate (fp) and *Mnx1*-expressing alleles in the pmn in comparison to expression of both genes in the ZPA, foregut (fg) and lung buds (l) of an E10.5 embryo assayed by RNA-FISH. Data for a biological replicate are in Extended Data Fig. 1b. Number of alleles scored [N] is shown below.

Here we investigate the co-activation of *Shh* and *Mnx1* by enhancers within the Shh TAD, using nascent RNA-FISH to determine whether expression of *Shh* and *Mnx1* alleles on the same chromosome can co-occur or are mutually exclusive, and DNA-FISH to examine the spatial context of such gene activations. We also explore the role of cohesin in activation of *Shh* and *Mnx1* driven by Shh enhancers in the Shh TAD. Our results have implications for models of enhancer-promoter communication and the role of spatial proximity, loop extrusion, and TAD boundaries, in developmental gene regulation.

## Results

### Mnx1 transcription is detected in some Shh-expressing tissues

*Shh* and *Mnx1* have very different zones of expression in the developing neural tube. *Shh* is expressed in the floorplate, regulated by enhancers located close to, and within, the *Shh* gene at the centromeric end of the TAD (Anderson et al., 2014). *Mnx1* in the adjacent TAD is expressed in the developing motor neurons of the neural tube, driven by enhancers just upsteam of the *Mnx1* promoter (Nagano et al., 2005) (Fig. 1a-c). *Shh* expression in the ZPA, at the posterior distal margin of the developing limb bud, is driven by the ZRS, 900kb upstream of Shh and at the opposite end of the *Shh* TAD (Lettice et al., 2003; Sagai et al., 2005). The ZRS is only 150kb from *Mnx1* in the adjacent TAD (Fig. 1a). Evidence for *Mnx1* expression in the developing mouse limb bud comes from quantitative RT-PCR (qRT-PCR) of limb buds and from low resolution in situ hybridisation (Williamson et al., 2019) and lacZ staining (Arber et al.,1999) of the developing embryo. To confirm that this expression were detected in the ZPA (Fig. 1d), with nascent *Mnx1* RNA detected for 9 - 20% of alleles, approximately half the levels for *Shh* and significantly lower than pre-motor neurons (Fig. 1e, h, Extended Data Fig. 1a, b, Extended Data Table 1). Detectable expression of both genes was minimal in the dorsal neural tube and locations adjacent to the ZPA in the posterior limb bud (Fig. 1c, d, Extended Data Fig. 1c). There were negligible levels of *Mnx1* transcription in the floorplate, or of *Shh* in pre-motor neurons of the neural tube, indicating that the enhancers proximal to each gene are unable to activate the other gene (Extended Data Fig. 1c, Extended Data Table 2). Together these data would be consistent with the ZRS being able to activate *Mnx1* in the ZPA despite the presence of an intervening TAD boundary.

We have previously shown that a 35kb deletion (35 del) of the TAD boundary separating ZRS from *Mnx1*, and including the first two exons of *Lmbr1* and 13 kb upstream of the *Lmbr1* promoter, leads to increased *Mnx1* expression in the ZPA as detected by in situ hybridisation to embryonic limb sections and qRT-PCR on dissected limb buds (Williamson et al., 2019). We confirmed a significant increase in transcription at *Mnx1* in the ZPA of 35 del homozygous embryos by RNA-FISH, with no apparent effects on *Shh* expression levels (Fig. 1e, Extended Data Fig. 1a, Extended Data Table 1).

A bit further into the Shh TAD from the ZRS the MACS1 enhancer, located in intron 8 of *Rnf32* (Fig. 1a), drives *Shh* expression in the epithelial linings of the laryngotracheal tube, stomach and lungs (Sagai et al., 2017). We therefore considered that *Mnx1* might also be responsive to this *Shh* enhancer. Indeed dual colour RNA FISH revealed transcription at *Mnx1*, along with the expected *Shh* expression, in the epithelium of the ventral foregut and the lung bud of E10.5 embryos (Fig. 1f, g). Although the proportion of active *Mnx1* alleles are approximately one third of those of *Shh* in these tissues (Fig.1h), the levels of *Mnx1* transcription in these epithelial cells, as well as in the ZPA, are significantly greater than in non-expressing *Shh* tissue and in the floorplate where *Shh* expression is driven by its proximal enhancers, over 1Mb away from Mnx1 (Extended Data Table 2). These data suggest that transcription from the *Mnx1* locus can be activated by *Shh* enhancers located at least a few 100s of kb away in the neighbouring TAD.

### Mnx1 and Shh can be transcribed simultaneously from the same allele

Conventional enhancer-promoter looping models predict that transcriptional bursts from two genes under the control of the same enhancer should not be coincident. However, in *Drosophila* there is evidence for co-ordinated transcriptional bursts from two reporter genes under the control of a single enhancer (Fukaya et al., 2016). Whether there could be coupled transcription from two endogenous genes driven by the same enhancer has not been explored.

We therefore used RNA-FISH to determine whether transcription from *Mnx1* and *Shh* originate from the same, or alternate, alleles. In the ZPA, the foregut, and the lung buds, the majority of *Mnx1* RNA-FISH signals are at alleles that show simultaneous signal for *Shh* nascent transcript from the same allele (closely apposed signals) (Fig. 2a, b and Extended Data Fig. 2a). In del 35 embryos, an even higher proportion of *Mnx1* transcribing alleles also transcribe *Shh* (Fig. 2b, Extended Data Fig. 2a, Extended Data Table 3.). These data suggest that both the ZRS and MACS1 enhancers are able to simultaneously activate transcription at two gene loci on the same chromosome.

**Figure 2.**
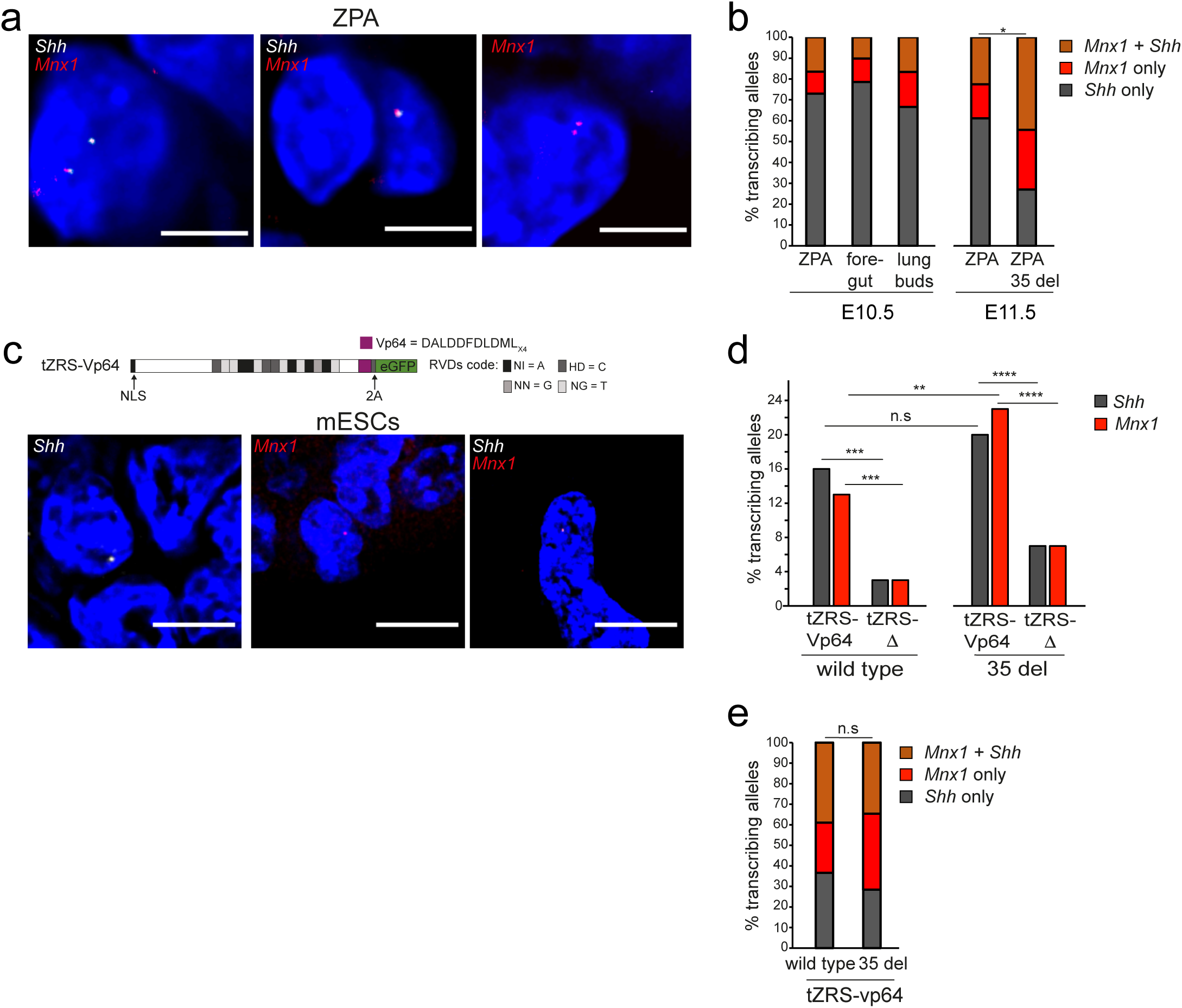
Adjacent RNA FISH signals signify co-activation at *Shh* and *Mnx1* by long-range enhancers. **a,** Representative images of ZPA nuclei from E10.5 embryos showing RNA-FISH signal at *Shh* (white) and *Mnx1* (red) at the same allele (left and centre panels), at *Shh* only (left panel upper allele) or at *Mnx1* only (right panel). Scale bars, 5 μm. **b,** Bar graph showing the % of active alleles transcribing at both *Shh* and *Mnx1* (brown), or at *Shh* (grey) or *Mnx1* (red) alone, in the ZPA, ventral foregut, and lung bud epithelial cells of a wild type E10.5 embryo (left) and in the ZPA of a wild type and 35 del E11.5 embryo (right). Co-activation at both genes versus activation of a single gene at expressing alleles of the wild type and 35 del E11.5 embryos was compared using a two-sided Fisher’s exact test; *, *P* ≤ 0.05 and > 0.01. Data from a biological replicate are in Extended Data Fig. 2a. Co-activation proportions in E11.5 wild type and 35 del cells and statistical data are in Extended Data Table 3. The proportion of transcribed alleles where either gene or both were detected and statistical analysis on the significance of co-activation by a single enhancer for E10.5 and E11.5 tissues are in Extended Data Table 4. **c,** Schematic of the TALE-Vp64 construct used to target the ZRS (tZRS-Vp64) enhancer (above). NLS, nuclear localization sequence; 2A, self-cleaving 2A peptide. Repeat variable diresidue (RVD) code is displayed alongside using the one-letter amino acid abbreviations. Equivalent TALE-Δ constructs lack the VP64 module. Below; Representative images of mESC nuclei showing RNA-FISHsignal for *Shh* (white) only, *Mnx1* (red) only, and both at the same allele. Scale bars, 5 μm. **d,** % of *Shh*- and *Mnx1*-transcribing alleles in wild type (left) and 35 del (right) mESCs activated from the ZRS targeted by either tZRS-Vp64 or tZRS-Δ. The data were compared using a two-sided Fisher’s exact test; n.s., not significant; **, *P* < 0.01; ***, *P* < 0.001; ****, *P* < 0.0001. Data from a biological replicate are in Extended Data Fig. 2b. Number of alleles scored, proportions transcribed, and statistical data are in Extended Data Table 5. **e,** As in (**b**) but for wild type and 35 del mESCs transfected with tZRS-Vp64. Data from a biological replicate are in Extended Data Fig. 2c. Statistical data comparing wild type and 35 del are in Extended Data Table 3. Statistical analysis of the frequency of co-activation by tZRS-Vp64 of *Shh* and *Mnx1* on the same allele vs different alleles in wild type and 35 del mESCs are in Extended Data Table 4.

To test whether there was an association between the transcription of pairs of genes on the same chromosome activated by the same enhancer, we considered cells where transcription was detected by RNA-FISH at exactly one copy of each gene. We classified these cells into two categories: those where there was transcription at both *Shh* and *Mnx1* on the same chromosome and those where transcription at each locus happened on different chromosomes. Replicates were merged due to the reduced number of cells suitable for this analysis. We then fitted generalized linear models with a binomial link function to assess both the level of significance (if any) and the direction of the association, i.e. whether alleles were preferentially expressed on the same or on different chromosomes. No tissue or cell type examined showed preferential transcription of *Shh* and *Mnx1* on different chromosomes, providing no evidence in support of an enhancer working exclusively on one gene at a time (Extended Data Table 4). In contrast, in the ZPA of E11.5 embryos, and in wild-type mESCs transfected with tZRS-Vp64, this analysis supported preferential simultaneous *Shh* and *Mnx1* transcription from the same chromosome (Extended Data Table 4).

ZRS and MACs1 are both relatively large and complex enhancers (>800bp) that bind a cocktail of different transcription factors in developing tissues to regulate *Shh* expression (Lettice et al., 2012; Lettice et al., 2017; Peluso et al.,2017;Sagai et al., 2017; Rankin et al.,2021; Huang et al., 2023). However, we have shown that transcription at *Shh* can also be induced by the binding of single synthetic transcription factors, to either the *Shh* promoter or to long-range *Shh* enhancers including the ZRS, in mouse embryonic stem cells (mESCs) (Benabdallah et al., 2019; Kane et al., 2022). We therefore set out to determine whether TAL effector (TALE) directed binding of Vp64 targeted to the ZRS could also induce transcription at *Mnx1* in mESCs. We found that, compared to a control TAL lacking fusion to Vp64 (tZRS-Δ), TAL-Vp64 targeted to the ZRS (tZRS-Vp64) could significantly induce transcription at both *Shh* and *Mnx1* in mESCs (Fig. 2c,d, Extended Data Fig. 2b). Whilst the deletion at the TAD boundary near ZRS in 35 del mutant mESCs had no significant effect on the ability of tZRS-Vp64 to activate transcription from *Shh*, it significantly enhanced transcription at *Mnx1* (Fig. 2d, Extended Data Fig. 2b, Extended Data Table 5). Therefore an enhancer (ZRS), activated by a single species of transcription factor (tZRS-Vp64) is able to activate transcription at a gene locus across a TAD boundary, and this is enhanced in the presence of a deletion encompassing that boundary. As in the limb bud (Fig. 2b), RNA-FISH indicated that tZRS-Vp64 can activate transcription simultaneously at both *Shh* and *Mnx1* on the same chromosome in mESCs (Fig. 2e, Extended Data Fig. 2c, Extended Data Table 4).

### Simultaneous transcription at Mnx1 and Shh occurs in the context of a compact chromosome conformation

To determine whether the ability of the ZRS to activate *Mnx1*, despite their location in separate TADs, is linked to spatial proximity of the genes and enhancers in the nucleus, we used DNA-FISH for *Shh*, *ZRS* and *Mnx1*, on developing limb bud tissue sections previously analysed by RNA-FISH (Fig. 3a and Extended Data Fig. 3a). We were therefore able to compare spatial distances between *Shh* and *Mnx1* and the ZRS enhancer in the ZPA of the distal posterior limb bud for alleles with no nascent transcription detected at either gene, or transcription detected at one gene or the other, or at both. Strikingly, transcription from *Mnx1* occurred in the context where the *Shh - Mnx1, Mnx1* - ZRS, and *Shh* - ZRS distances were all small. *Mnx1*-*Shh* and *Mnx*-ZRS distances were larger at alleles where no transcription from *Mnx1* was detected (Fig. 3b, Extended Data Fig. 3b, Extended Data Table 6). Taking as a cut-off 350 nm, encompassing the upper quartile of inter-probe distances between *Shh*-ZRS at all ZPA *Shh*-transcribing alleles (Extended Fig. 3c), >50% of alleles in the limb-bud show *Shh* and ZRS in close proximity even when no *Shh* transcription is detected, but this proportion increases even further at *Shh* transcribing alleles (Fig. 3c). In contrast, <25% of alleles where no transcription at *Mnx1* is detected have *Mnx*-ZRS or *Mnx1*-*Shh* distances <350nm, but this rises very significantly to 50-70% at *Mnx1* transcribing alleles (Extended Data Table 7). This is a consequence of ZRS-driven activation, not *Mnx1* transcription per se. In the nuclei of pre-motor neurons, where *Mnx1* expression is driven from its own proximal enhancers (Fig. 1a), *Mnx*-ZRS and *Mnx1*-*Shh* distances are not different between *Mnx1* expressing and non-expressing alleles (Fig. 3d-f, Extended Data Fig. 3d, Extended Data Table 7).

**Figure 3.**
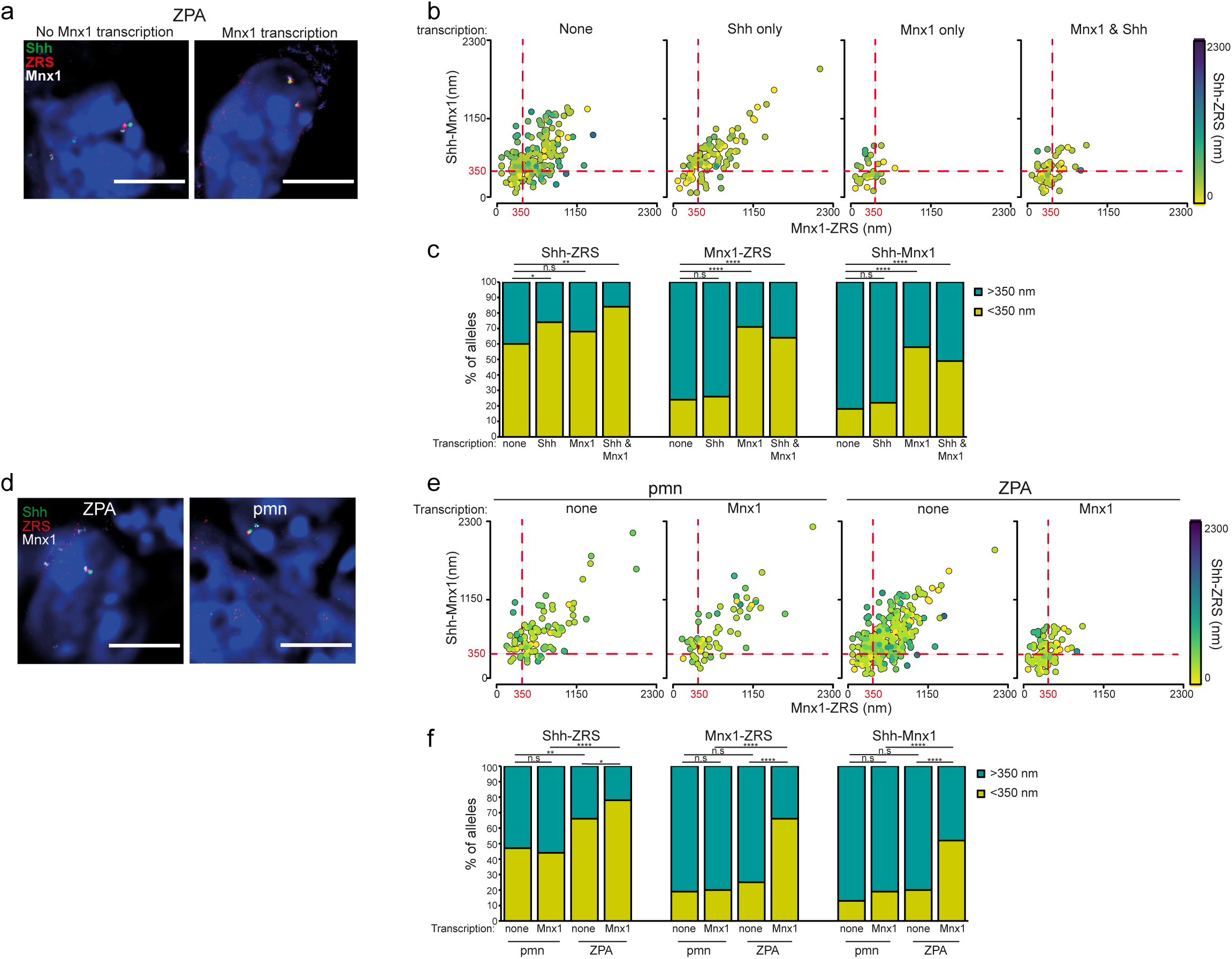
Spatial proximity of *Mnx1* to ZRS and *Shh* is optimal for limb bud ZPA transcription. **a,** Representative images of ZPA nuclei from E10.5 embryos showing DNA FISH signal for *Shh* (green), ZRS (red) and *Mnx1* (white) at *Mnx1*-non-transcribing (left) and transcribing (right) alleles. Scale bars, 5 μm. **b,** Scatter plots showing interprobe distances between each of the two probe pairs indicated on *x* and *y* axes with the separation between the third pair indicated by the colour (in the colour bar) in ZPA cells at non-, *Shh*-, *Mnx1*- and *Mnx1* & *Shh*-transcribing alleles. Dashed red lines indicates alleles where *Shh*-*Mnx1* or *Mnx1*-ZRS inter-probe distances are < 350nm. **c,** Bar plots providing categorical analysis of the spatial relationship of *Shh*, ZRS and *Mnx1* in ZPA cells at non-, *Shh*-, *Mnx1*- and *Mnx1* & *Shh*-transcribing alleles. Categories are: < 350 nm apart, the upper (75%) quartile distance of all *Shh*-transcribing alleles (Extended Data Fig. 3b); > 350 nm. Differences between non-expressing and expressing alleles identified using Fisher’s Exact test; n.s., not significant; *, *P* ≤ 0.05 and > 0.01; **, *P* ≤ 0.01; ****, *P* ≤ 0.0001. Data from two biological replicates is combined in **b** and **c**. Median and interquartile data for each of the two biological replicates are in Extended Data Fig. 3c. Statistical analysis of the significance of spatial proximity at transcribing versus non-transcribing alleles and the proportion of interprobe distances < 350 nm are in Extended Data Tables 6 and 7. **d,** Representative images of a ZPA (left) and neural tube pre-motor neuron (pmn; right) nuclei showing DNA FISH signal for *Shh* (green), ZRS (red) and *Mnx1* (white). Scale bars, 5 μm. **e,** As in (**b**) but for pmn and ZPA cells at non-*Mnx1*- and all *Mnx1*-transcribing alleles. **f,** As in (**c**) but in pmn and ZPA cells at non-*Mnx1*- and *Mnx1*-transcribing alleles. Differences between non-expressing and expressing alleles identified using Fisher’s Exact test; n.s., not significant; *, *P* ≤ 0.05 and > 0.01; **, *P* ≤ 0.01; ****, *P* ≤ 0.0001. All data from both biological replicates combined. Median and interquartile data for each of the two biological replicates are in Extended Data Fig. 3d. Statistical analysis of the significance of spatial proximity at pmn *Mnx1*-transcribing alleles versus pmn non-transcribing alleles and *Mnx1* transcribing and non-transcribing alleles in ZPA cells and the proportion of interprobe distances < 350 nm are in Extended Data Tables 6 and 7.

### Cohesin facilitates activation of Mnx1 across the Shh TAD boundary

It has been shown that cohesin, and likely cohesin-mediated loop extrusion, is required for enhancers to activate transcription at target genes across large (several hundreds of kb) genomic distances (Calderon et al., 2022; Kane et al., 2022; Rinzema et al., 2022) but not for more short-range interactions. Indeed, we have previously shown that the ability of the synthetic transcription factor tZRS-Vp64 to activate transcription from *Shh* in mESCs depends on cohesin (SCC1/RAD21) but not on CTCF (Kane et al., 2022). We therefore set out to examine whether the ability of tZRS-Vp64 to activate transcription at *Mnx1* across the intervening TAD boundary, was influenced by the degradation of CTCF or SCC1. We transfected tZRS-Vp64 into mESCs engineered to contain AID-degron tagged CTCF or SCC1 (Nora et al., 2017; Rhodes et al. 2020) and then degraded these tagged proteins by the addition of auxin as previously described (Kane et al., 2022). Consistent with deletion of CTCF sites in the mouse (Williamson et al., 2019) and with previous studies (Kane et al., 2022), CTCF degradation resulted in no significant change in the ability of tZRS-Vp64 to induce transcription at *Shh*. Similarly, tZRS-Vp64 induced transcription at *Mnx1* was not significantly affected by CTCF degradation (Fig. 4a, Extended Data Fig. 4a, Extended Data Table 8). As previously shown (Kane et al., 2022), acute cohesin depletion very significantly blunted the ability of tZRS-Vp64 to activate transcription at *Shh*. Notably, it also reduced the ability of tZRS-Vp64 to induce transcription at *Mnx1*, but to a lesser extent than for *Shh* (Fig. 4a, Extended Data Fig. 4a, Extended Data Table 8). Therefore, in the absence of cohesin, tZRS-Vp64 more efficiently induced transcription at *Mnx1* than at *Shh*, likely a reflection of the smaller linear genomic separation of ZRS and *Mnx1*.

**Figure 4.**
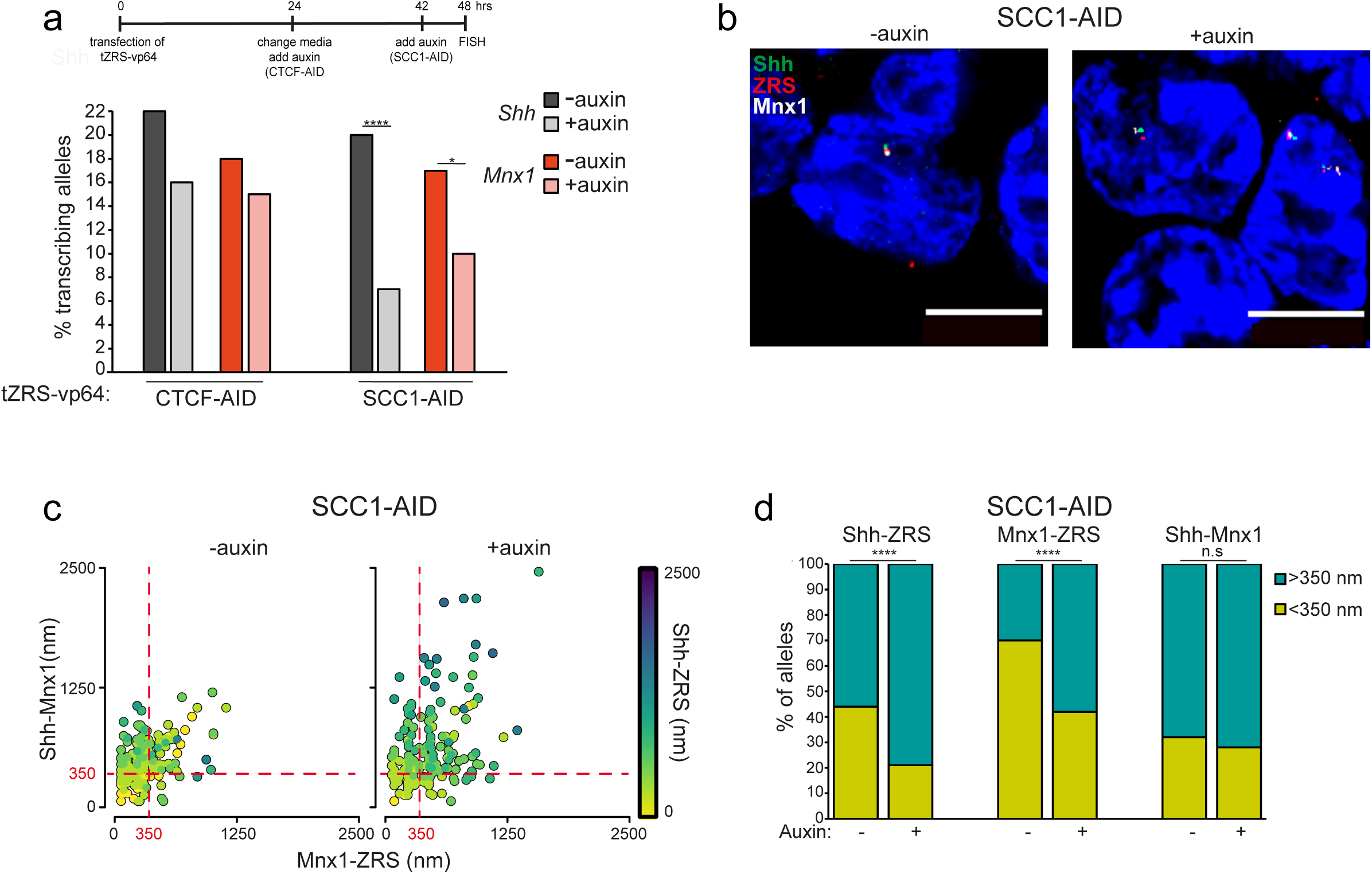
Cohesin optimises *Mnx1* activation across the *Shh* TAD boundary. **a,** Top; timecourse of TALE transfection and auxin treatment. Below; % of *Shh*- and *Mnx1*-transcribing alleles in TALE-transfected CTCF-AID cells (left) and SCC1-AID cells (right) either untreated (- auxin) or treated with 24 hours (CTCF-AID) or 6 hours (SCC1-AID) of auxin (+ auxin) mESCs activated from the ZRS targeted by tZRS-Vp64. Data were compared using a two-sided Fisher’s exact test; n.s., not significant; *, *P* ≤ 0.05 and > 0.01; ****, *P* < 0.0001. Data from a biological replicate are in Extended Data Fig. 4a. Values for number of alleles scored and statistical evaluation are summarised in Extended Data Table 8. **b,** Representative images of untreated (- auxin) and treated (+ auxin) SCC1-AID nuclei showing DNA-FISH probe signal for *Shh* (green), ZRS (red) and *Mnx1* (white). Scale bars, 5 μm. **c,** Scatter plots showing interprobe distances between each of the two probe pairs indicated on *x* and *y* axes with the separation between the third pair indicated by the colour (in the colour bar) in SCC1-AID cells untreated (- auxin) and treated (+ auxin). Dashed red lines indicates alleles where *Shh*-*Mnx1* or *Mnx1*-ZRS inter-probe distances are <350 nm. **d,** Bar plots providing categorical analysis of the spatial relationship of *Shh*, ZRS and *Mnx1* in SCC1-AID mESCs - or + auxin. Categories are: < 350 nm and >350 nm. Differences between - & + auxin cells were identified using Fisher’s Exact test; n.s., not significant; ****, *P* ≤ 0.0001. Data from two biological replicates is combined in **c** and **d**. Median and interquartile data for each of the two biological replicates are in Extended Data Fig. S4c. Statistical analysis of the significance of spatial proximity at transcribing versus non-transcribing alleles and the proportion of interprobe distances <350 nm are in Extended Data Tables 9 and 10.

We previously showed by DNA-FISH that significant decompaction of the *Shh* TAD occurs in the absence of cohesin (Kane et al., 2022). Therefore, as expected and consistent with our previous data, we found that the spatial distances between ZRS-*Shh* significantly increased following the acute depletion of cohesin (SCC1/RAD21). Interestingly, ZRS-*Mnx1* distances also significantly increased following cohesin depletion (Fig. 4b, c, Extended Data Fig. 4b, Extended Data Table 9). The proportion of alleles with closely apposed ZRS-*Mnx1* loci (<350nm) also decreased when cohesin was degraded (Fig. 4d & Extended Data Table 10). Together, these data indicate that it is likely to be cohesin-mediated loop extrusion running through the TAD boundary that is maintaining ZRS-Mnx1 proximity in the nucleus.

We noted that, whilst there was an increased incidence of a few alleles with very large distances between *Shh* and Mnx1 in the absence of Rad21/Scc1, the average *Shh* – *Mnx1* distances and the proportion of alleles with closely apposed *Shh*-*Mnx1* loci (<350nm) were not significantly affected by cohesin loss despite the large genomic distance (∼1Mb) separating these two genes (Fig. 4, Extended Data Fig. 4, Extended Data Tables 9 and 10). We consider that this is likely due to both *Shh* and *Mnx1*genes being targets of the polycomb complexes in mESCs and forming strong polycomb-mediated loops in mESCs (Fig. 4c,d & Extended Data Fig. 4b,c) (Boyle et al., 2020; Williamson et al., 2023).

## Discussion

How the hundreds of thousands of enhancers in the mammalian genome activate their target genes, sometimes over intervening genes and large genomic distances, without inadvertently activating other nearby genes, remains an unanswered question. Alignment of the large regulatory domains of developmental genes within TADs implicates 3D genome organsation and cohesin-mediated loop extrusion in both facilitating and restraining enhancer action (Symmons et al., 2014; 2016, Andrey and Mundlos, 2017). In particular, it has been proposed that intact TAD boundaries help to prevent enhancers in one TAD activating genes in neighbouring TADs, with genetic, and epigenetic, disruption of TAD boundaries giving rise to ectopic gene activation by enhancers and disease phenotypes (Flavahan et al., 2016; 2019; Hnisz et al., 2015; Lupianez et al., 2015; 2016; Tsujimura et al., 2015).

However, experiments to test this hypothesis have not reached a consensus. In many cases, small deletions of TAD boundaries do not seem to affect developmental gene regulation or result in mutant phenotypes (Despang et al., 2019; Paliou et al., 2019; Williamson et al., 2019). The insertion of ectopic CTCF sites can influence some enhancers but not others (Chakraborty et al., 2023) but, on the other hand, removal of CTCF sites at TAD boundaries can result in ectopic gene activation in cancer cell lines (Kim et al., 2024) and have been associated with Mendelian disease (Baudic et al., 2024). The phenotypic consequence of deleting regions containing CTCF sites close to long-range enhancers can also differ between mouse and human (Ushiki et al., 2021).

The *Shh* locus is an exemplar for a large regulatory domain with >15 enhancers distributed across an ∼900kb TAD of the human or mouse genomes, including in the introns of other genes. Aberrant *Shh* expression or developmental phenotypes are not seen in mice with deletion of individual CTCF sites at the *Shh* TAD boundaries (Paliou et al., 2019; Williamson et al., 2019). For example, deletion of the CTCF sites at the TAD boundary closest to *Shh* (Fig.1a) does not result in ectopic *Shh* expression detectable by in situ hybridisation driven by the *En2* and *Cnpy1* enhancers (Li Song and Joyner, 2000; Hirate and Okamoto, 2006; Williamson et al., 2019). However, we previously noted that the other end of the *Shh* TAD, close to the location of the ZRS Shh limb enhancer, mRNA for *Mnx1* – a gene located in the adjacent TAD - can be detected by in situ hybridisation in the *Shh* expressing ZPA of developing limb buds (Williamson et al., 2019).

Here, we use RNA FISH, detecting nascent transcription, to confirm transcription at *Shh* or *Mnx1* in nuclei of the same cells of the ZPA. Nascent transcription at *Mnx1* and *Shh* could be detected simultaneously from the same allele, often at a higher frequency than expected by chance. Therefore, the *Shh* ZRS enhancer can simultaneously activate transcription at two genes and across an intact, but porous, TAD boundary. The simultaneous transcription at two endogenous mammalian loci driven by an intervening single enhancer is consistent with the observation of simultaneous transcription of two reporter genes by a shared enhancer in Drosophila embryos (Fukaya et al., 2016).

*Mnx1* transcription in the ZPA is further enhanced by a 35kb deletion encompassing the TAD boundary separating the ZRS from *Mnx1*, suggesting either that the *Shh* TAD boundary near the promoter of *Lmbr1* has some ability to insulate *Mnx1* from the influence of the ZRS, that is due to more than just CTCF binding, or that reducing the genomic separation between ZRS and *Mnx1* from 150 to 115kb has a significant effect. Of note, and consistent with our observations, a lacZ reporter transgene (SBLac936) inserted upstream of the *Lmbr1* promoter, just beyond the Shh TAD boundary, shows evidence of weak expression driven by ZRS in the limb, and by *Mnx1* enhancers in the the neural tube (Anderson et al., 2014).

It has been suggested that sequences within the ZRS recruit specific transcription factors that endow the enhancer with the ability to act over large genomic distances in the developing limb (Lettice et al., 2014; Bower et al., 2024). However, we also detect nascent transcription from *Mnx1* in the *Shh* expressing portions of the developing ventral foregut and the lung bud of E10.5 embryos, an activity that is driven by the *Shh* MACS1 enhancer, located a further 100kb into the Shh TAD from ZRS (Sagai et al., 2017) and therefore able to induce transcription at *Mnx1* across a TAD boundary from a distance of >260 kb (Fig. 1a). We also demonstrate simultaneous activation of nascent transcripts at *Mnx1* and *Shh* using a synthetic transcription factor (tZRS-Vp64) targeted to the ZRS in mESCs. This suggests that activation of nascent transcription over long genomic distances and across a TAD boundary is not dependent on the recruitment of a specific cocktail of transcription factors.

Combining RNA- and DNA-FISH we show that transcription activation of *Mnx1* from the ZRS in the ZPA of developing limb buds occurs in the context of a compact chromatin conformation, where Mnx1 is close (<350nm) to both *Shh* and the ZRS. This is not the case when *Mnx1* transcription is being driven by its own proximal enhancer in pre-motor neurons. Together with the observation of simultaneous transcription at *Mnx1* and *Shh,* which does not seem compatible with classical enhancer-promoter looping models (Lim and Levine, 2021), this result is consistent with a model in which enhancers nucleate the formation of transcription hubs which can activate transcription at genes coming into their sphere of influence (Fig. 5). We have previously shown that within the *Shh* TAD cohesin, but not CTCF, is required for a synthetic transcription factor to activate transcription at *Shh* from a large genomic distance (100s of kb) (Kane et al., 2022). In the absence of cohesin, and presumably cohesin-mediated loop-extrusion, decompaction of the TAD was observed, suggesting that one of the functions of loop-extrusion is to compact large chromatin domains and to thereby facilitate the localisation of target genes within the sphere of influence of enhancers located distant in the linear genome. Here we show this also applies across a TAD boundary – the ability of a transcription factor targeted to the ZRS to activate transcription at *Mnx1* is attenuated when cohesin is degraded (Fig. 5). Interestingly, this is to a lesser extent than the influence of cohesin degradation on activation at *Shh*, such that transcription is now detected at a higher proportion of *Mnx1* alleles, compared with *Shh*. This likely reflects the shorter genomic distance (150kb) separating *Mnx1* from ZRS, compared with *Shh* (850kb).

**Figure 5.**
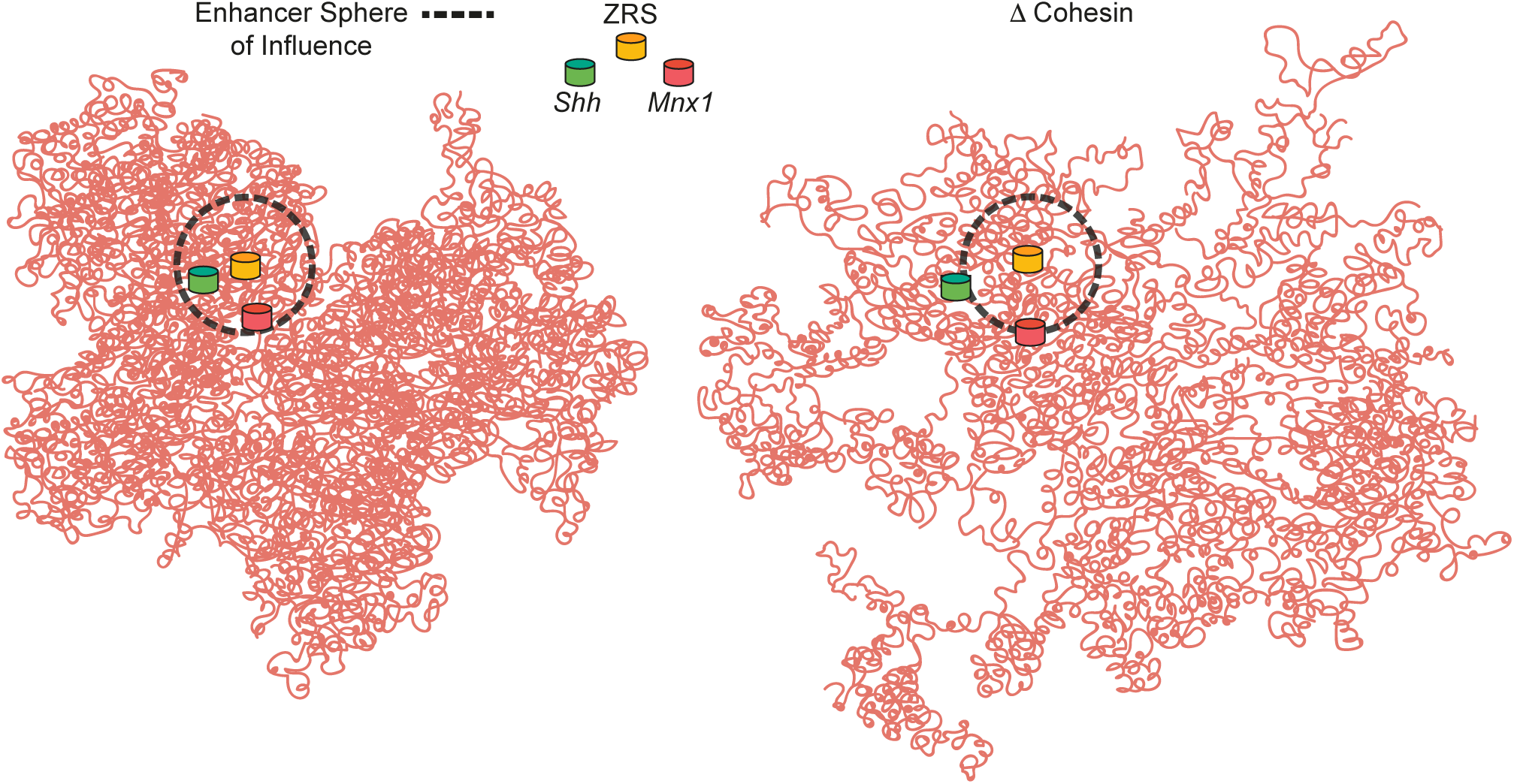
Cohesin and the ZRS sphere of influence. Cohesin-mediated chromatin compaction of the *Shh* TAD enables regulatory signals from the ZRS enhancer to reach and activate *Shh* >850 kb away, and also to enable some activation at *Mnx1,* much closer to ZRS genomically, but located the other side of a TAD boundary (left). Acute loss of cohesin results in a decompaction of the chromatin domain, putting *Shh* beyond the reach of signals from the ZRS and considerably reducing the opportunity for *Mnx1* to be within the enhancer’s ‘sphere of influence’ (right).

Consistent with our previous genetic analysis of CTCF sites at the *Shh* locus in cells and in mice (Williamson et al., 2019), we detect no significant increase in ZRS-driven transcription at *Mnx1* when CTCF is degraded, even though the latter results in a loss of insulation at this TAD boundary as assayed by Hi-C (Nora et al., 2017; Kane al., 2022). In cancers, loss of CTCF binding due to altered DNA methylation has been suggested to result in ectopic enhancer-driven expression of cancer driver genes (Flavahan et al., 2016; 2019; Kim et al., 2024). It will therefore be important to better understand the different factors that contribute to functional TAD boundaries at different sites in the mammalian genome, and in different cellular contexts.

Our findings suggest that not all TAD boundaries act as absolute barriers to enhancer function, and that ‘ectopic’ activation of genes from enhancers in adjacent TADs depends, to some extent, on the action of cohesin – and presumably loop extrusion, resulting in spatial proximity of the enhancer and ‘ectopic’ target gene. In the situation studied here, of the ectopic activation of *Mnx1* by enhancers in the *Shh* TAD, this appears to have no discernable biological function. The extent of such ectopic activation across TAD boundaries in the mammalian genome remains to be explored, but it is interesting to speculate that it may provide a framework on which evolution can act to drive new patterns of gene expression.

## Methods

### Mouse strains and embryo sectioining

Embryos were collected from timed matings of wildtype and the 35 del (Williamson et al, 2019) mouse lines and were fixed, embedded, sectioned and processed for FISH as previously described (Morey et al., 2007; Williamson et al., 2019), except that sections were cut at 8 μm.

All mouse work has been ethical approved by the University of Edinburgh Animal Welfare and Ethics Review board and is conducted under the authority of Home Office Licences.

### Cell culture and treatments

Mouse embryonic stem cells (mESCs) used were wild type E14 (parental line of the CTCF-AID cells provided by Elphege Nora), CTCF-AID (Nora et al., 2017) and SCC1-AID (Rhodes et al., 2020) and the homozygous 35 del line (Williamson et al., 2019).

Feeder-free mESCs were cultured and transfected as previously described (Kane et al., 2022). Briefly 1.5 x 10^6^ mESCs were transfected with 14.5 μg of TALE plasmid and 26 μL Lipofectamine 3000 Reagent (Invitrogen L3000015) and seeded onto 0.1% gelatin-coated 10 cm dishes containing autoclaved SuperFrost Plus Adhesion glass slides. Fresh media was added after 24 hours (h). After 48 h of transfection, slides were washed, fixed in 4% paraformaldehyde (pFa) and permeabilised in 70% ethanol at 4°C for minimum of 24 h (up to one week). For auxin-inducible protein degradation, cells from half of each TALE transfection were treated with 500 µM auxin and half left untreated for an internal control. Auxin was added to the CTCF-AID 24 h after transfection and left for a further 24 h, while the SCC1-AID cells received auxin 42 h after transfection and were treated for a further 6 h.

### RNA-FISH

Custom Stellaris® RNA FISH Probes were designed against *Shh* nascent mRNA (a pool of 48 22-mers probes designed to (NCBI37/mm9: chr5:28,787,847-28,793,741) and *Mnx1* nascent mRNA (a pool of 46 22-mers probes designed to (NCBI37/mm9 chr5:29,800,772-29,804,270) by utilizing the Stellaris® RNA FISH Probe Designer (Biosearch Technologies, Inc., Petaluma, CA) (Biosearch Technologies, Inc.), following the manufacturer’s instructions available online at www.biosearchtech.com/stellarisprotocols and as previously described (Williamson et al, 2019).

### 3D DNA-FISH

Following RNA FISH, slides were re-probed by DNA FISH. After the removal of coverslips, slides were briefly washed in PBS and then for 5 mins in 2xSSC at 75°C (80°C for mESCs) followed by denaturation in 70% formamide/2xSSC at 75°C for 20 minutes (80°C for 50 minutes for mESCs) before a series of alcohol washes (70% (ice-cold), 90% and 100%). 160-240 ng of biotin- and digoxigenin-labeled and red-dUTP-labeled (Alexa Fluor™ 594-5-dUTP, Invitrogen) or Green496-dUTP-labeled (Enzo Life Sciences) fosmid probes (Extended Data Table 11) were used per slide, with 16-24 µg of mouse Cot1 DNA (Invitrogen) and 10 µg salmon sperm DNA. EtOH was added and the probe air dried. Hybridisation mix containing deionised formamide, 20 x SSC, 50% dextran sulphate and Tween 20 was added to the probes for ∼1h at room temperature. The hybridisation mix containing the probes was added to the slides and incubated overnight at 37°C. Following a series of washes in 2X SSC (45°C) and 0.1X SSC (60°C) slides were blocked in blocking buffer (4 x SSC, 5% Marvel) for 5 min. The following antibody dilutions were made: fluorescein anti-dig FAB fragments (Roche cat. no. 11207741910) 1:20, fluorescein anti-sheep 1:100 (Vector, cat. no. FI-6000)/ streptavadin Cy5 1:10 (Amersham, cat. no. PA45001, lot 17037668), biotinylated anti-avidin (Vector, cat. no. BA-0300, lot ZF-0415) 1:100, and streptavidin Cy5 1:10. Slides were incubated with antibody in a humidified chamber at 37°C for 30 -60 min in the following order with 4X SSC/0.1% Tween 20 washes in between: fluorescein anti-dig, fluorescein anti-sheep/streptavidin Cy5, biotinylated anti-avidin, streptavidin Cy5. Slides were treated with 1:1000 dilution of DAPI (stock 50ug/ml) for 5min before mounting in Vectashield.

### Image acquisition and deconvolution

Slides from RNA and DNA FISH using fosmid derived probes were imaged on an epifluorescence microscope as previously described (Boyle et al., 2020). Step size for z stacks was set to 0.2 µm. Hardware control and image capture were performed using Nikon Nis-Elements software (Nikon) and images were deconvolved using a calculated PSF with the constrained iterative algorithm in Volocity (PerkinElmer). RNA FISH signal quantification was carried out using the quantitation module of Volocity (PerkinElmer). Number of expressing alleles were calculated by segmenting the hybridization signals and scoring each nucleus as containing 0, 1 or 2 RNA signals. DNA FISH measurements were made using the quantitation module of Volocity (PerkinElmer) and only alleles with single probe signals were analysed to eliminate the possibility of measuring sister chromatids.

### Statistical analysis

DNA-FISH interprobe distance data sets were compared using the two-tailed Mann-Whitney U test, a nonparametric test that compares two unpaired groups. Differences in DNA-FISH data sets comparing categorical distributions, differences in the proportion of *Shh* and *Mnx1* transcribing alleles in wild type and 35kb deletion embryos and mESCs transfected with either tZRS-Vp64 or tZRS-Δ, and comparisons between levels of transcribing alleles in SCC1-AID mESCs ± auxin transfected with tZRS-Vp64, were measured using Fisher’s Exact Test. These statistical analyses were performed using GraphPad Prism 9.4.1 software (Mann-Whitney, T-test) or online GraphPad 2×2 contingency table www.graphpad.com/quickcalcs/contingency1/ (Fisher’s).

The association between the transcription of *Shh* and *Mnx1* regulated by the same enhancer was done by linear modelling with binomial link function. Significance of association was determined using the Chi-square Test. The analysis was carried out in R (version 4.4.1) using the built-in function ’glm’.

## Author contributions

I.W., R.E.H. W.A.B and L.A.L. conceived the project. R.E.H. W.A.B and L.A.L supervised the project. I.W. designed and performed experiments, and analysed data. K.A.G. performed some RNA-FISH. H.B. performed the binomial tests. I.W. conducted all other statistical tests. I.W., W.A.B and L.A.L. prepared the figures and wrote the paper.

## Acknowledgements

We thank the University of Edinburgh BVS, the Advanced Imaging Resource at the Institute of Genetics and Cancer and the Genome Engineering Core at the MRC Human Genetics Unit for their technical support. This work has made use of the resources provided by the Edinburgh Compute and Data Facility (ECDF) (http://www.ecdf.ed.ac.uk/). We are grateful to Elphege Nora and Rob Klose for their gift of their CTCF- and Rad21/SCC1-AID degron mESC lines.

Work in the group of W.A.B. is supported by MRC University Unit grant MC_UU_00035/7. K.A.G. was supported by an MRC PhD studentship. Funding sources were not involved in study design, data collection, data interpretation, or the decision to submit the work for publication.

## Extended Data Figure Legends

**Extended Data Figure 1. Related to Figure 1.**
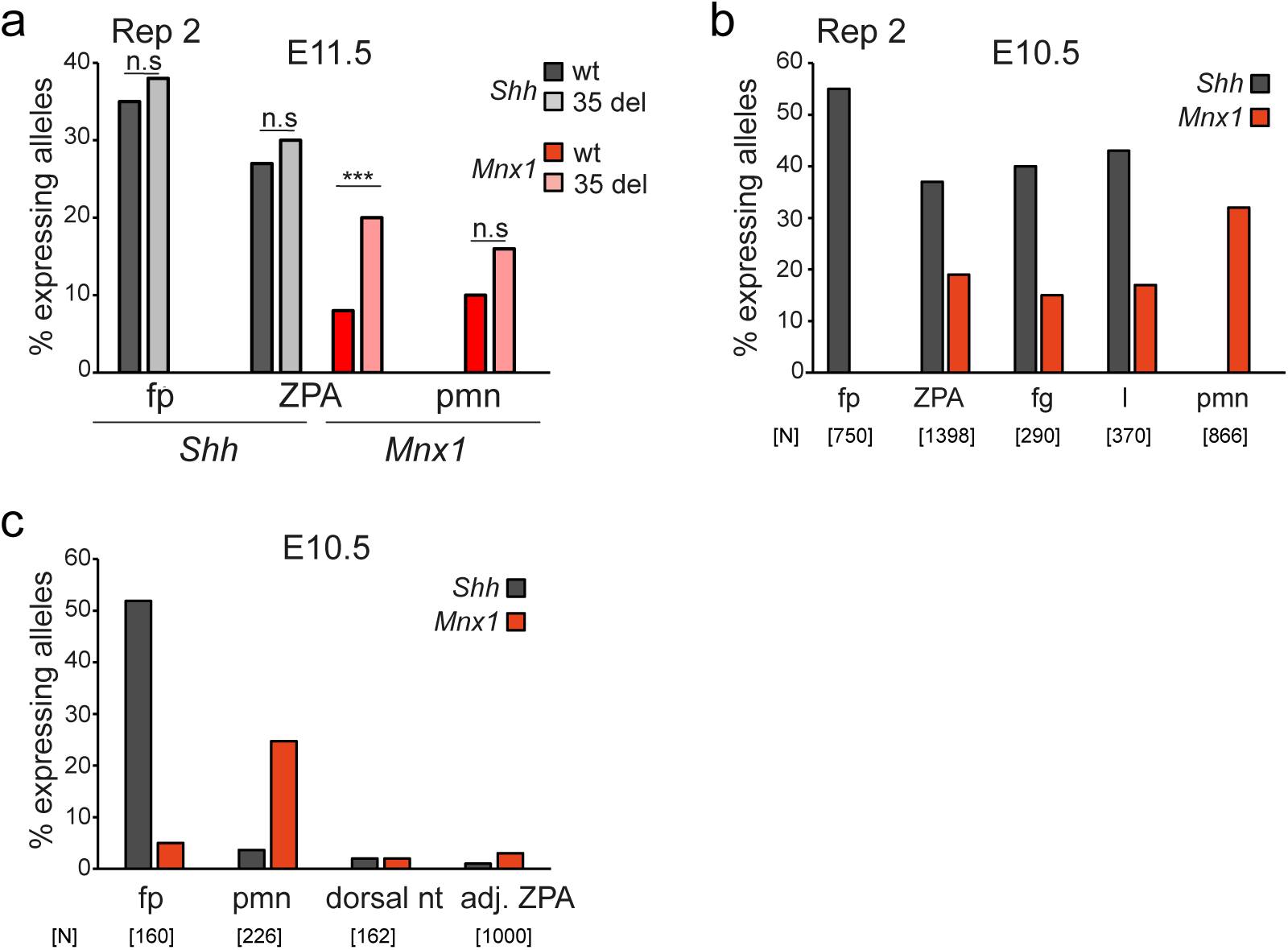
**a,** Percentage of alleles with *Shh* and *Mnx1* RNA FISH signal in wild type and 35 kb deletion (35 del) E11.5 mouse embryos in tissues of the floorplate (fp), ZPA and pre-motor neurons (pmn). The data were compared using a two-sided Fisher’s exact test; n.s., not significant; ***, *P* < 0.001. Data are for a biological replicate of Figure 1e. Number of alleles scored, and proportions transcribed, and statistical data are in Extended Data Table 1. **b,** Percentage of *Shh*-expressing alleles in the floorplate and *Mnx1*-expressing alleles in the pmn in comparison to the expression of both genes in the ZPA, foregut (fg) and lung buds (l) of an E10.5 embryo. Data are for a biological replicate of Fig. 1h. Number of alleles scored [N] are shown below. **c,** The percentage of alleles with *Shh* and *Mnx1* RNA-FISH signal in the fp, pmn, dorsal neural tube (nt) and locations adjacent (adj.) to the ZPA of wild type E10.5 mouse embryos. Number of alleles [N] scored are shown below.

**Extended Data Figure 2. Related to Figure 2.**
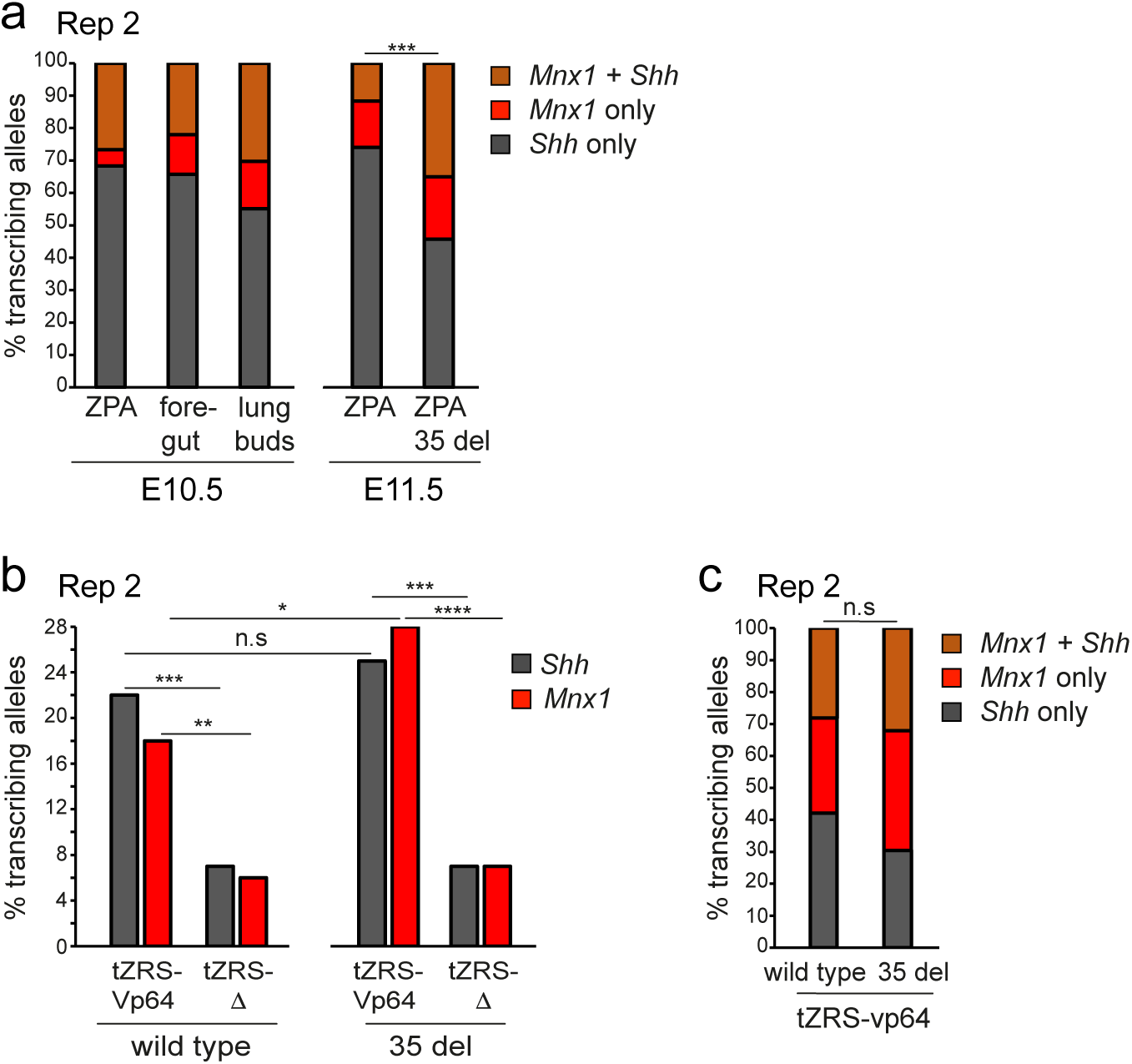
**a,** Bar graph showing the proportion of active alleles transcribing at both *Shh* and *Mnx1* (brown), or at *Shh* (grey) or *Mnx1* (red) alone, in the ZPA, ventral foregut, and lung bud epithelial cells of a wild type E10.5 embryo (left) and in the ZPA of a wild type and 35 kb deletion (35 del) E11.5 embryo (right). Co-activation at both genes versus activation of a single gene at expressing alleles of the wild type and 35 del E11.5 embryos was compared using a two-sided Fisher’s exact test; ***, *P* < 0.001. Data are a biological replicate for Fig. 2b. Co-activation proportions in E11.5 wild type and 35 del cells and statistical data are in Extended Data Table 3. Statistical analysis of the frequency of co-activation of Shh and Mnx1 on the same allele vs different alleles in E10.5 and E11.5 tissues are in Extended Data Table 4. **b,** % of *Shh*- and *Mnx1*-transcribing alleles in wild type (left) and 35 del (right) mESCs activated from the ZRS targeted by either tZRS-Vp64 or tZRS-Δ. The data were compared using a two-sided Fisher’s exact test; n.s., not significant; *, *P* ≤ 0.05 and > 0.01, **, *P* < 0.01; ***, *P* < 0.001; ****, *P* < 0.0001. Data are from a biological replicate for Fig. 2d. Number of alleles scored, proportions transcribed, and statistical data are in Extended Data Table 5. **c,** % of *Shh*/*Mnx1* transcriptional co-activation to activation at either *Shh* or *Mnx1* alone for transcribing alleles in wild type and 35 del mESCs transfected with tZRS-Vp64. Data are for a biological replicate of Fig. 2e. Co-activation proportions in wild type and 35 deletion cells are in Extended Data Table 3. Statistical analysis on the significance of co-activation by the ZRS enhancer for E14 wild type and 35 deletion mESCs are in Extended Data Table 4.

**Extended Data 3. Related to Figure 3.**
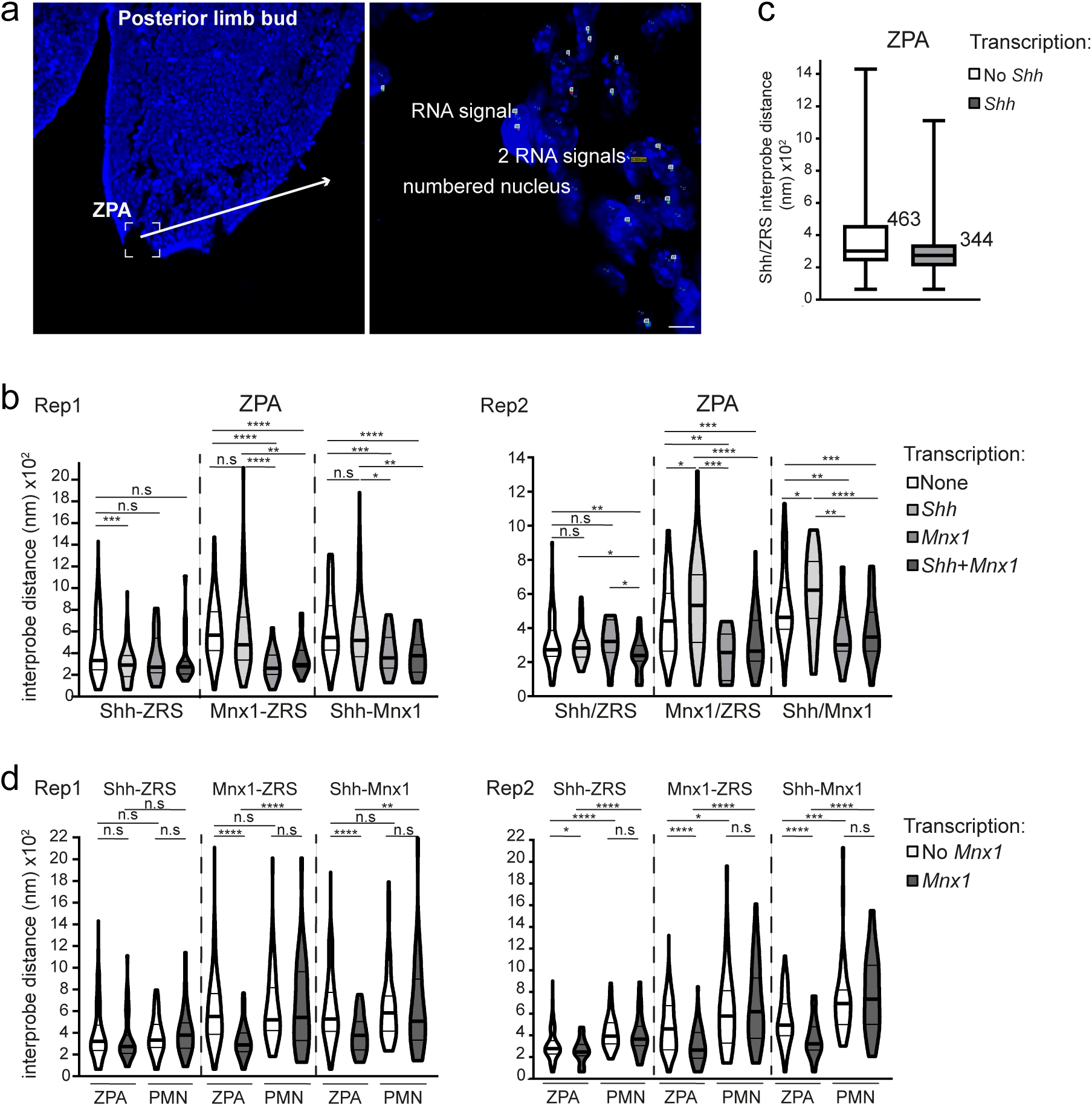
**a,** Representative images of E10.5 limb bud tissue section (left) that had been analysed by RNA-FISH (right) prior to DNA-FISH. **b,** Violin plots showing the distribution of interprobe distances (nm) between *Shh*-ZRS, *Mnx1*-ZRS, *Shh*-*Mnx1* probes in ZPA cells from two biological replicates at non-, *Shh*-, *Mnx1*- and *Mnx1* & *Shh*-transcribing alleles. **c,** Box plots showing the distribution of *Shh*-ZRS interprobe distances (nm) at *Shh* non-transcribing and transcribing alleles in ZPA cells from the two biological replicates combined. The upper quartile distances for non-transcribing and transcribing tissues are shown, the latter value determining the <350 nm category for optimal enhancer spatial proximity for Fig. 3. **d,** As in (**b**) but for pre-motor neuron (pmn) and ZPA cells at non-*Mnx1*- and all *Mnx1*-transcribing alleles. The data were compared using a two-sided Mann-Whitney U-test; n.s., not significant; *, *P* ≤ 0.05 and > 0.01; **, *P* < 0.01; ***, *P* < 0.001; ****, p<0.0001. Values for number of alleles scored, median and inter-quartile distances, and statistical evaluation are summarised in Extended Data Table 6.

**Extended Data 4. Related to Figure 4.**
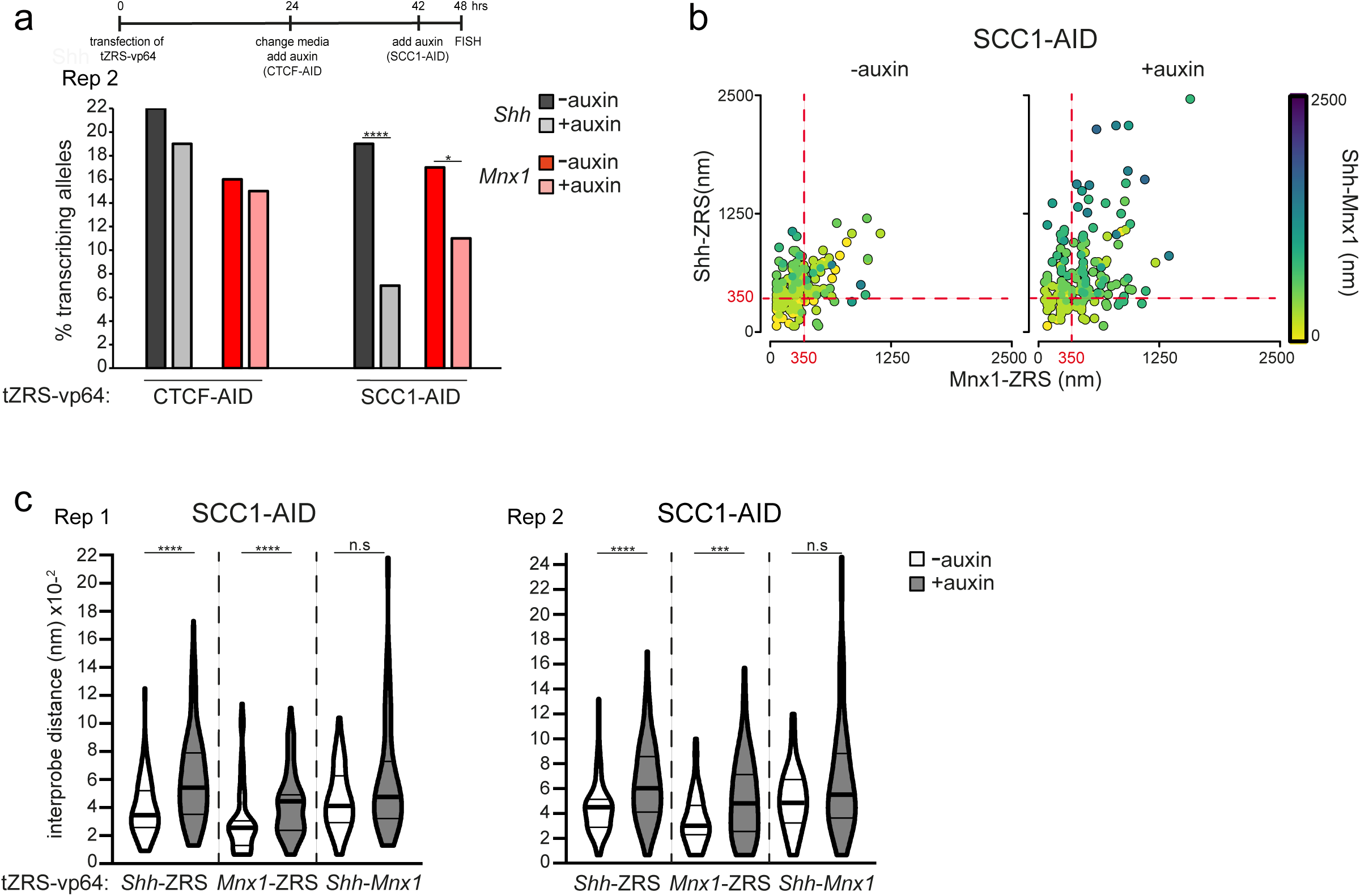
**a,** Top; timecourse of TALE transfection and auxin treatment. Below; % of *Shh*- and *Mnx1*-transcribing alleles in TALE-transfected CTCF-AID cells (left) and SCC1-AID cells (right) either untreated (- auxin) or treated with 24 hours (CTCF-AID) or 6 hours (SCC1-AID) of auxin (+ auxin) in mESCs activated from the ZRS targeted by tZRS-Vp64. The data were compared using a two-sided Fisher’s exact test; n.s., not significant; *, *P* ≤ 0.05 and > 0.01; ****, *P* < 0.0001. Values for number of alleles scored and statistical evaluation are summarised in Extended Data Table 8. Data are from a biological replicate of Fig. 4a. **b,** Scatter plots showing interprobe distances between each of the two probe pairs indicated on *x* and *y* axes with the separation between the third pair indicated by the colour (in the colour bar) in SCC1-AID cells - & + auxin. Dashed red lines indicates alleles where *Shh*-*Mnx1* or *Mnx1*-ZRS inter-probe distances are <350 nm. Same data as in Fig. 4c but with *Shh*-*Mnx1* rather than *Shh*-ZRS interprobe distances colour-coded. **c,** Violin plots showing the distribution of interprobe distances (nm) between *Shh*-ZRS, *Mnx1*-ZRS, *Shh*-*Mnx1* probes in tZRS-Vp64-transfected SCC1-AID cells - or + auxin. Data from two biological replicates are shown. Data were compared using a two-sided Mann-Whitney U-test; n.s., not significant; ***, *P* < 0.001; ****, *P* < 0.0001. Values for number of alleles scored, median and inter-quartile distances, and statistical evaluation are summarised in Extended Data Table 9.

**Extended Data Table 1.**
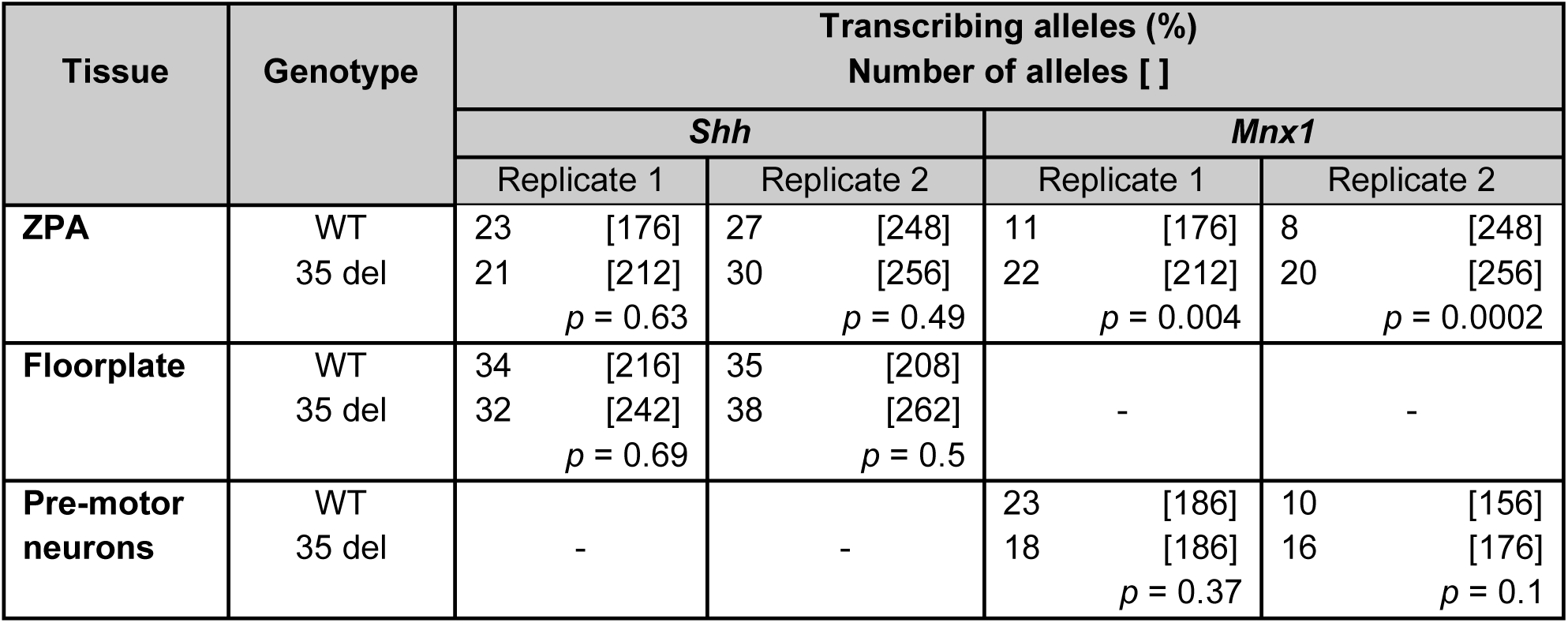
The proportion of *Shh* and *Mnx1* transcribing alleles in wild type and 35kb deletion E11.5 limb bud ZPA and neural tube tissues. Statistical analysis of data for Fig. 1e & Extended Data Figure 1a. Table shows the percent of alleles transcribing at *Shh* and *Mnx1* as assayed by RNA FISH signal in wild type (WT) and 35 kb deletion (35 del) E11.5 mouse embryos in tissues of the floorplate, ZPA and pre-motor neurons. Number of alleles scored is indicated in square brackets. *p* values are from two-sided Fisher’s exact test comparing data from wild-type and 35 del embryos.

**Extended Data Table 2.** Significance of the frequency of *Mnx1* and *Shh* transcription activated by enhancers of the other gene. Statistical analysis of data for Extended data Figs. 1b & c. Comparison of *Mnx1* transcription in *Shh*-expressing tissues of E10.5 embryos, regulated by either *Shh* proximal or distal enhancers, and comparison of transcription of both genes in non-expressing tissue with tissue expressing the other gene. *Shh* and *Mnx1* non-expressing tissues: dorsal neural tube and limb bud adjacent to the ZPA. *Mnx1*-expressing tissue (to identify significant *Shh* activation by *Mnx1* enhancers): pre-motor neurons (proximal *Mnx1* enhancers). *Shh*-expressing tissues: floorplate (proximal *Shh* enhancers), and ZPA, foregut, lung buds (distal *Shh* enhancers). The distances (kb) between Mnx1 and the relevant Shh enhancer is shown. N/A = not applicable. *p*-values from Fisher’s Exact Tests.

| E10.5 *Shh*-exp tissue<br>(*Shh* Non-exp [ ]) | Distance *Shh* enhancer to *Mnx1* (kb) | Transcribing alleles (%) |  |
| --- | --- | --- | --- |
|  |  | *Mnx1* | *Shh* |
| [Dorsal neural tube] | N/A | 2 | 2 |
| [adj. ZPA] | N/A | 3 | 1 |
| [Pre-motor neurons] | N/A | 25 | 5<br>($p = 0.41$) |
| **Floorplate** | >1000kb | 5<br>($p = 0.14$) | 52 |
| **Foregut** | 250kb | 15<br>($p < 0.0001$) | 40 |
| **Lung buds** | 250kb | 17<br>($p < 0.0001$) | 43 |
| **ZPA** | 150kb | 20<br>($p < 0.0001$) | 37 |

**Extended DataTable 3.**
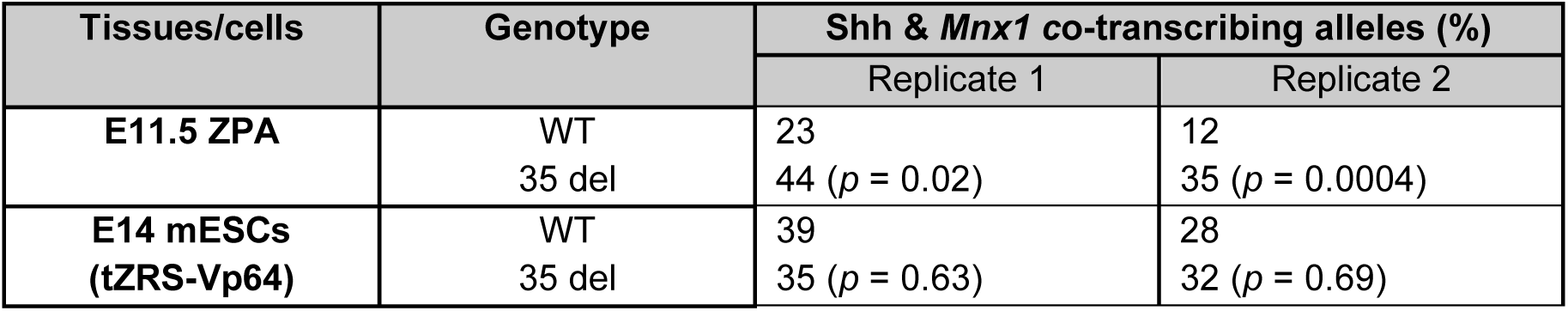
Comparison of *Shh* and *Mnx1* co-transcription in wild type and 35kb deletion ZPA and mESCs. Statistical analysis of data for Figs. 2b,e & Extended Data Figs. 2a,c. Table shows the percent of alleles transcribing at *Shh* and *Mnx1* as assayed by RNA-FISH signal in wild type (WT) and 35 kb deletion (35 del) in the ZPA of E11.5 mouse embryos and in mESCs transfected with tZRS-Vp64. Values are proportion of total transcribed alleles. *p*-values from Fisher’s Exact Tests.

**Extended Data Table 4.**
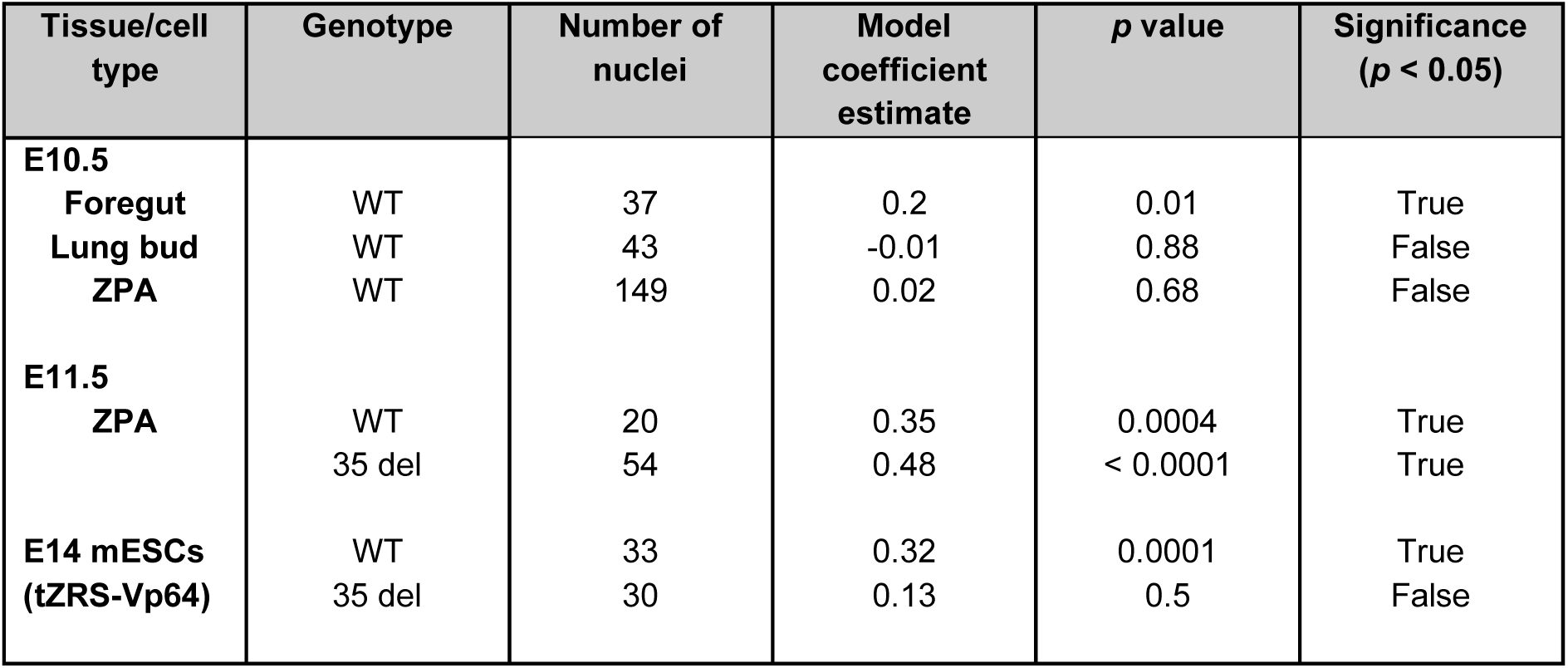
Binomial test comparison of the frequency of *Shh* and *Mnx1* co-transcription. Data associated with Figs. 2b,e & Extended Data Fig. 2a,c. Analysis in E10.5 tissues from wild-type (WT) animals and in E11.5 tissue and E14 mESCs for both WT and 35kb del genotypes. E14 mESCs were transfected with tZRS-Vp64. Only nuclei containing one *Shh* and one *Mnx1* signal were analyzed using fitted generalized linear models with a binomial link function to determine if there were more nuclei where both genes are being transcribed on the same chromosome or where they are being transcribed on separate chromosomes. A coefficient estimate value close to 0 indicates equal numbers of co-transcribed and individually transcribed alleles, a positive value indicates preferential co-transcription and a negative value indicates preferential individual transcription. *P*-values from Chi-square Tests.

**Extended Data Table 5.**
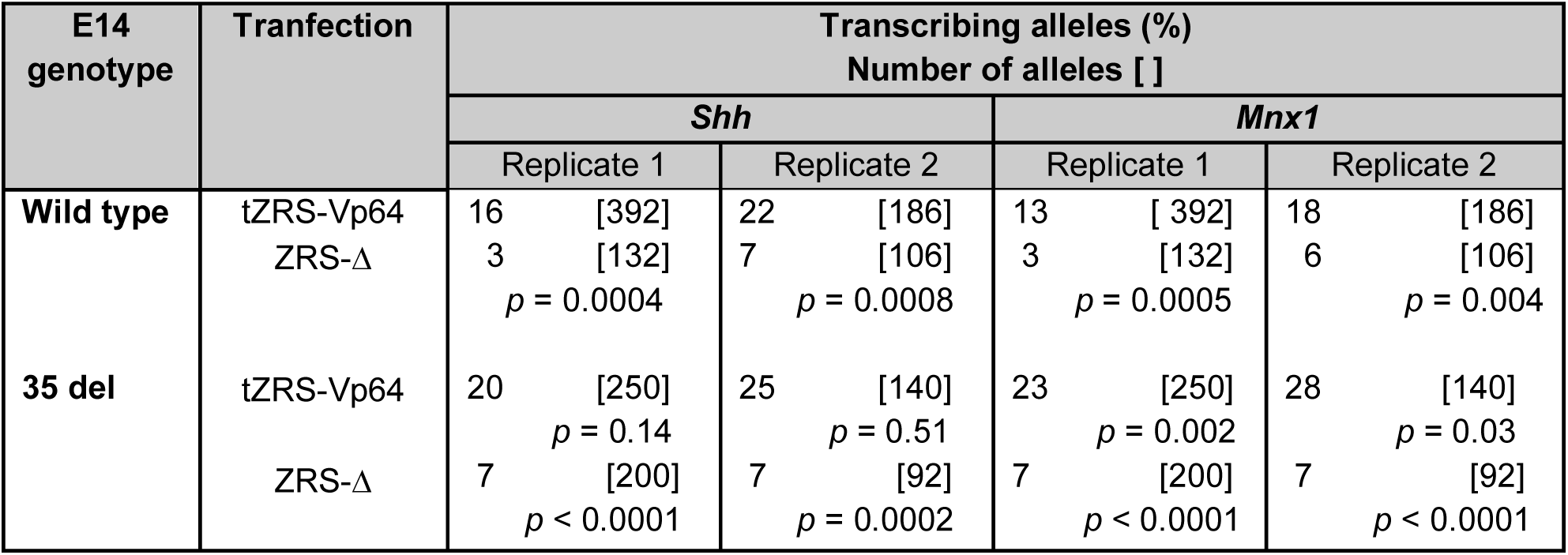
Proportion of *Shh* and *Mnx1* transcribing alleles in E14 wild type and 35kb deletion mESCs transfected with either tZRS-Vp64 or tZRS-Δ. Statistical analysis of data for Fig. 2d & Extended Data Fig. 2b. *p*-values from Fisher’s Exact Tests. tZRS-Δ values were compared with those from tZRS-Vp64 for the same cell type. 35 del cells transfected with tZRS-Vp64 were compared with wild type cells transfected with tZRS-Vp64.

**Extended Data Table 6.**
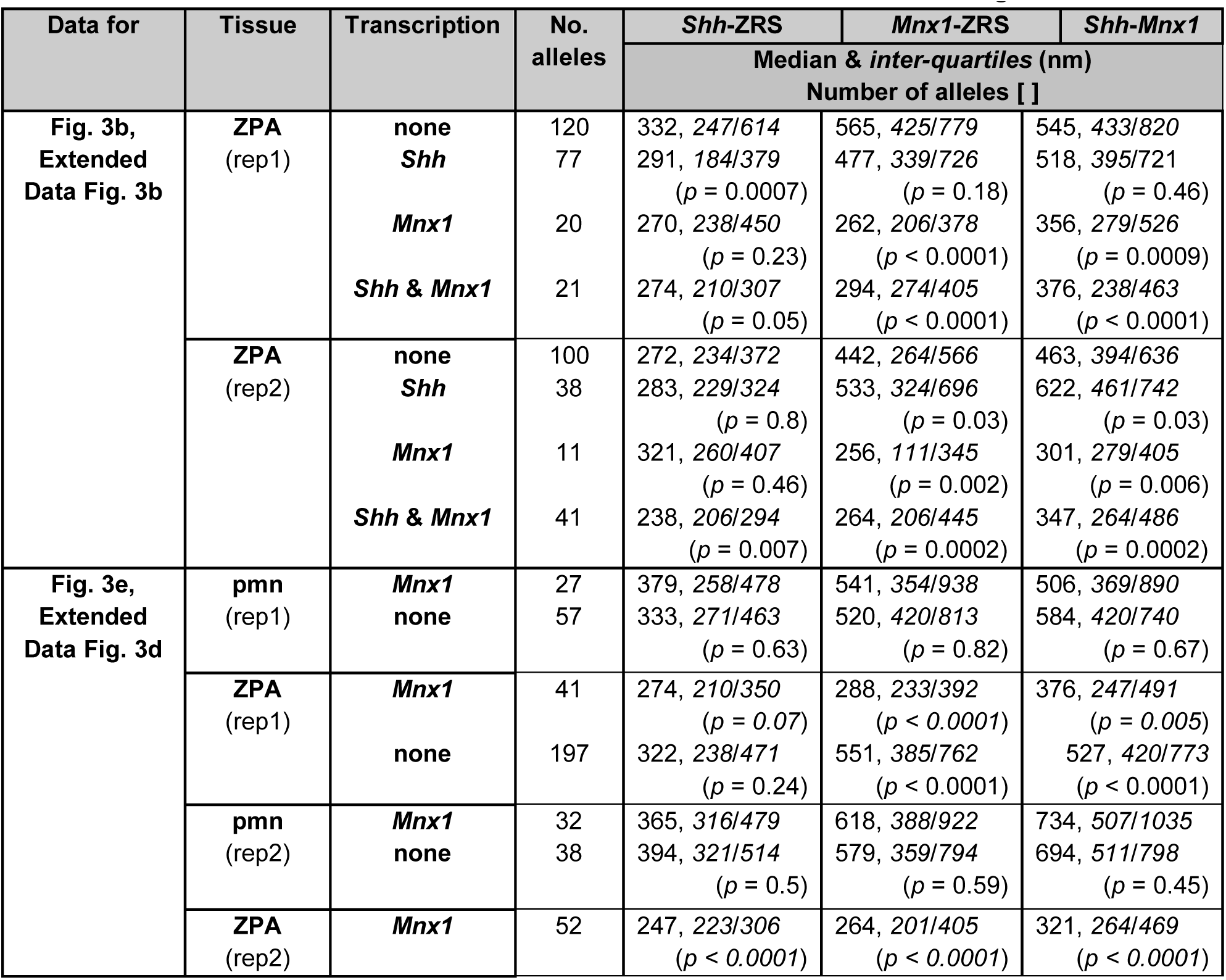
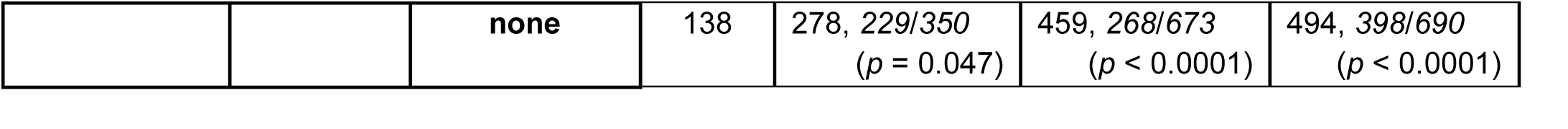
Distribution of *Shh*-ZRS, *Mnx1*-ZRS, *Shh*-*Mnx1* interprobe distances in ZPA cells at non-, *Shh*-, *Mnx1*- and *Mnx1* & *Shh*-transcribing alleles. Statistical analysis of DNA-FISH *Shh-ZRS*, *Mnx1-ZRS* and *Shh-Mnx1* inter-probe distance data for Fig. 3b, e. and Extended Data Fig. 3b, d. Data are from two biological replicates with Number of alleles scored shown. Median distances (nm) are shown together with inter-quartile distances in italics. *p*-values from Mann-Whitney U Tests. For Fig. 3b/ED Fig. 3b data *p* values are in comparison to alleles not transcribing either *Shh* or *Mnx1*. For Fig3e/ED Fig. 3d data *p* values *Mnx1*-transcribing and non-transcribing values were compared for each tissue (ZPA or pre-motor neuron (pmn)) and *Mnx1*-transcribing values in pmn tissues were compared with ZPA *Mnx1*-transcribing values.

**Extended Data Table 7.** Categorical analysis of the spatial relationship of *Shh*, ZRS and *Mnx1* in ZPA. Categorical analysis of the inter-probe distances between *Shh*-ZRS, *Mnx1-ZRS* and *Shh-Mnx1* in ZPA and PMN cells at non-, *Shh*-, *Mnx1*- and *Shh* & *Mnx1*-transcribing alleles (Fig. 3c) and at non- and *Mnx1*-transcribing alleles (Fig. 3f). Statistical analysis of data for Figs. 3c & f with replicate data sets combined. *p*-values from Fisher’s Exact Tests are for comparison to non-expressing alleles for Fig.3c data, and to *Mnx1* expressing alleles for data in Fig 3f.

| Data for | Tissue | Transcription | *Shh*-ZRS | *Mnx1*-ZRS | *Shh*-*Mnx1* |
| --- | --- | --- | --- | --- | --- |
|  |  |  | **Alleles < 350nm apart (%)** |  |  |
| **Fig. 3c** | **ZPA** | **none** | 60 | 24 | 18 |
|  |  | *Shh* | 74 (*p* = 0.02) | 26 (*p* = 0.67) | 22 (*p* = 0.45) |
|  |  | *Mnx1* | 68 (*p* = 0.44) | 71 (*p* < 0.0001) | 58 (*p* < 0.0001) |
|  |  | *Shh* & *Mnx1* | 84 (*p* = 0.001) | 64 (*p* < 0.0001) | 49 (*p* < 0.0001) |
| **Fig. 3f** | **PMN** | *Mnx1* | 47 | 19 | 13 |
|  |  | None | 44 (*p* = 0.74) | 20 (*p* = 0.84) | 19 (*p* = 0.36) |
|  | **ZPA** | *Mnx1* | 78 (*p* < 0.0001) | 66 (*p* < 0.0001) | 52 (*p* < 0.0001) |
|  |  | None | 66 (*p* = 0.03) | 25 (*p* < 0.0001) | 20 (*p* < 0.0001) |

**Extended Data Table 8.** Proportion of *Shh* and *Mnx1* transcribing alleles in E14 CTCF-AID and SCC1-AID - or + auxin mESCs transfected with tZRS-Vp64. Statistical analysis of data for Figs. 4a & Extended Data 4a examining the consequence of CTCF or SCC1 ablation (using auxin induced degradation) on the proportion of alleles transcribing at *Shh* or *Mnx1* in mESCs transfected with tZRS-Vp64.*p*-values from Fisher’s Exact Tests compare – vs + auxin data for two biological replicates. Number of alleles scored indicated in square brackets.

| E14 mESCs | Transcribing alleles (%) |  |  |  |
| --- | --- | --- | --- | --- |
|  | Number of alleles [ ] |  |  |  |
|  | *Shh* |  | *Mnx1* |  |
|  | Rep 1 | Rep 2 | Rep 1 | Rep 2 |
| **CTCF-AID** |  |  |  |  |
| - auxin | 22 [238] | 22 [134] | 18 | 16 |
| + auxin | 16 [300]<br>(*p* = 0.12) | 19 [132]<br>(*p* = 0.55) | 15<br>(*p* = 0.29) | 15<br>(*p* = 0.87) |
| **SCC1-AID** |  |  |  |  |
| - auxin | 20 [200] | 19 [218] | 17 | 17 |
| + auxin | 7 [290]<br>(*p* < 0.0001) | 7 [254]<br>(*p* < 0.0001) | 10<br>(*p* = 0.03) | 11<br>(*p* = 0.04) |

**Extended Data Table 9.** Distribution of distances between *Shh*-ZRS, *Mnx1*-ZRS, *Shh - Mnx1* DNA-FISH signals between in E14 SCC1-AID - or + auxin mESCs transfected with tZRS-Vp64. Statistical analysis of DNA-FISH *Shh-ZRS*, *Mnx1-ZRS* and *Shh-Mnx1* inter-probe distance data for Fig. 4c. and Extended Data Fig. 4b,c. Data are from two biological replicates with Number of alleles scored are indicated. Median distances (nm) are shown together with inter-quartile distances in italics. *p*-values from Mann-Whitney U Tests comparison data for – vs + auxin.

| SCC1-AID | No. of alleles | Median & *inter-quartiles* (nm) |  |  |
| --- | --- | --- | --- | --- |
|  |  | *Shh*-ZRS | *Mnx1*-ZRS | *Shh*-*Mnx1* |
| Rep1 |  |  |  |  |
| - auxin | 78 | 345, 258/520 | 255, 133/300 | 411, 295/616 |
| + auxin | 90 | 542, 351/786<br>(*p* < 0.0001) | 455, 238/485<br>(*p* < 0.0001) | 476, 323/720<br>(*p* = 0.05) |
| Rep2 |  |  |  |  |
| - auxin | 87 | 449, 288/514 | 300, 229/461 | 485, 334/668 |
| + auxin | 75 | 603, 419/857<br>(*p* < 0.0001) | 481, 259/710<br>(*p* = 0.0005) | 551, 370/861<br>(*p* = 0.09) |

**Extended Data Table 10.**
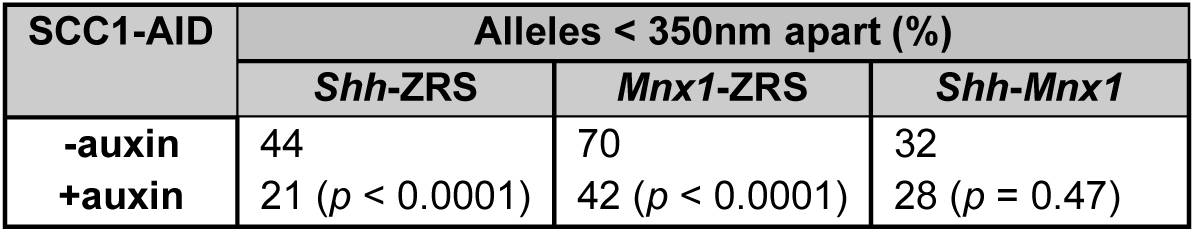
Categorical analysis of the spatial relationship of *Shh*, ZRS and *Mnx1* in E14 SCC1-AID - & + auxin mESCs. Statistical analysis of data for Figs. 4c & d. *p*-values from Fisher’s Exact Tests compare data for - vs + auxin..

**Extended Data Table 11.**
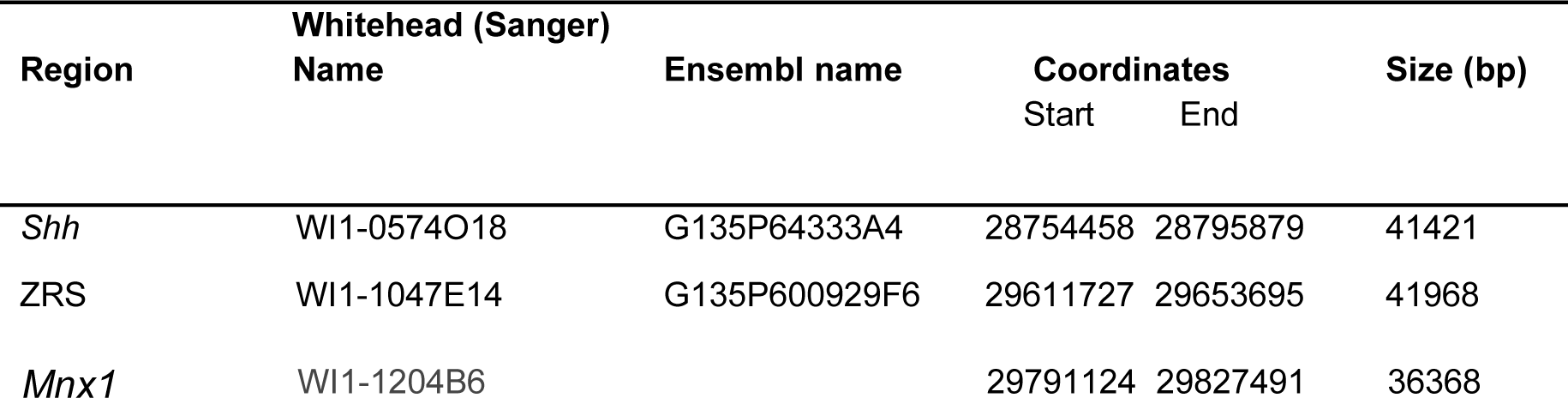
Fosmid Probes used in this study. Names are Ensembl (r 45) (http://jun2007.archive.ensembl.org/Mus_musculus/index.html). Mouse genome assembly number: NCBI m37.

